# Proteogenomic profiling of soft tissue leiomyosarcoma reveals distinct molecular subtypes with divergent outcomes and therapeutic vulnerabilities

**DOI:** 10.1101/2025.11.19.689365

**Authors:** Atsushi Tanaka, Makiko Ogawa, Yusuke Otani, Ronald C. Hendrickson, Zhuoning Li, Narasimhan P. Agaram, David S. Klimstra, Julia Y. Wang, Wenyi Wei, Michael H. A. Roehrl

## Abstract

Soft tissue leiomyosarcoma (STLMS) is an aggressive malignancy for which robust molecular subclassification and mechanism-based therapeutic strategies still remain limited. We performed integrative proteogenomic analyses of primary and metastatic STLMS to define subtype-associated molecular programs. Joint analysis of the proteome and phosphoproteome identified 3 biologically distinct subtypes. P1 was characterized by relative genomic stability, low proliferative activity, and enrichment of FGFR2- and PDK-associated signaling. In contrast, P2 and P3 showed greater chromosomal instability and more aggressive clinical behavior, but with distinct molecular features. Notably, P2 was associated with inflammatory and RTK-RAS pathway programs, activation of CDK-AURKA/B-mTOR-ERK kinase networks, IGF1R/PDGFRA alterations, and the poorest outcomes. On the other hand, P3 showed strong cell cycle and DNA repair programs, elevated NCOR1 expression, and increased expression of nonhomologous end joining components, including PARP1. Homologous recombination deficiency analyses distinguished HRD-low P1 from HRD-high P2/P3, and paired analyses suggested increased HRD-related features in metastatic lesions within P3. Immune profiling identified an immune-hot yet potentially suppressive state in P2, marked by higher LGALS9 expression and M2-like macrophage infiltration. To support clinical translation, we developed a tissue microarray-based immunohistochemical classifier that enabled surrogate assignment of proteome-defined subtypes in an independent cohort and showed recurrence-free survival differences across inferred subtypes. These findings together establish a proteogenomic framework for STLMS heterogeneity and nominate subtype-associated biological vulnerabilities for future translational and clinical investigation.

## Introduction

Soft tissue sarcomas (STS) are a diverse group of mesenchymal malignancies that include over 80 histological subtypes^1^. Leiomyosarcoma (LMS) accounts for 10-20% of all STS^2^. Soft tissue leiomyosarcoma (STLMS), i.e., extra-uterine LMS, develops mainly in the retroperitoneum, abdomen/pelvis, trunk, and extremities, with the first two being more common^3^. Clinical management of localized disease largely depends on anatomical site and tumor grade^4,5^. Typical STLMS shows spindle-shaped cells that are set in long intersecting fascicles, diffuse hypercellularity with high-grade histology, and the expression of one or more myogenic marker proteins. Despite multidisciplinary management, recurrence and distant metastasis remain major causes of mortality in STLMS. Half of patients experience tumor relapse after surgery^6–8^. Patients with advanced or metastatic STLMS have poor outcomes and limited treatment options^9,10^. Hence, improved biologic stratification may help clarify disease heterogeneity and identify clinically relevant subsets of STLMS.

Several genomic and epigenomic studies of pan-STS reported enrichment of PIK3/AKT pathway alterations and deletions of tumor suppressor genes, including *TP53* (9% deep and 60% shallow deletions), *RB1* (14% deep, 78% shallow), and *PTEN* (13% deep, 68% shallow)^11–13^. Multi-omic analyses showed that LMS is molecularly distinct from other STS, and that STLMS differs from uterine LMS in hormone response and DNA damage response pathways. Integrated clustering of STLMS based on WES, epigenome, and whole transcriptome data revealed two subtypes: one with short survival and frequent PI3K/AKT/MTOR signaling pathway alterations, the other with longer survival and minimal PI3K/AKT/MTOR alterations. While MTOR inhibitors like everolimus or temsirolimus show some clinical efficacy in LMS^14,15^, their efficacy is reduced by indirect AKT upregulation.

Proteins execute complex cellular and biochemical functions and are therefore important therapeutic targets in cancer^16^. Because mRNA abundance does not consistently predict protein levels^17,18^, proteomics provides complementary insight into tumor biology beyond nucleic acid-based approaches. Two recent pan-soft tissue sarcoma proteomic studies by Burns et al.^19^ and Tang et al.^20^ included leiomyosarcoma cases, but both were based on formalin-fixed, paraffin-embedded specimens and included LMS, including uterine leiomyosarcoma, rather than a cohort restricted to soft tissue LMS. Burns et al. performed global proteomic profiling and identified three LMS proteomic subtypes, whereas Tang et al. incorporated proteomic and phosphoproteomic analyses in a pan-sarcoma setting. However, a clearly matched soft tissue LMS-focused multi-omic cohort integrating DNA sequencing, transcriptome, proteome, and phosphoproteome has remained lacking. In addition, prior studies did not systematically analyze metastatic lesions, particularly paired primary-metastatic samples, which limited insight into proteogenomic changes associated with metastatic progression. Importantly, the use of FFPE material is a major constraint for deep proteomic profiling and an even greater limitation for phosphoproteomic analysis, because phosphorylation signals are particularly vulnerable to pre-analytical and fixation-related effects.

Here, we performed an integrative proteogenomic analysis of freshly frozen STLMS specimens with matched phosphoproteomic profiling. We analyzed 72 primary and metastatic STLMS samples, including 18 paired primary-metastatic lesions, together with matched DNA sequencing and transcriptomic data in a subset of cases, and incorporated an independent validation cohort of 219 STLMS FFPE samples as tissue microarrays. By incorporating phosphoproteomic information into subtype discovery, our study was designed to capture signaling pathway activity more directly, since phosphorylation is a key regulatory layer in tumor progression that is not fully reflected by DNA, RNA, or total protein abundance alone.

## Results

### Overview of study cohorts

We analyzed two independent STLMS cohorts with complementary purposes. Cohort 1 comprised 72 frozen primary and metastatic STLMS samples and served as the discovery cohort for integrative proteogenomic analyses, including global proteomics, phosphoproteomics, targeted DNA sequencing, and transcriptomic profiling in a subset of samples. Cohort 2 comprised 219 formalin-fixed, paraffin-embedded STLMS cases assembled into tissue microarrays and served as an orthogonal validation cohort for immunohistochemical evaluation of selected markers. We also used the TCGA soft tissue leiomyosarcoma cohort as an external multi-omic comparison dataset, as it provides broadly accessible molecular and clinical data. Although two additional sarcoma proteomic datasets have recently been reported^19,20^, limited access to accompanying genomic information and detailed sarcoma subtype annotation restricted their use for external comparison in this study. Unless otherwise specified, subtype discovery was performed in cohort 1, immunohistochemical validation was performed in cohort 2, and external comparison was performed using TCGA.

### Unsupervised clustering of proteome identifies 3 proteomic subtypes with clinical relevance

We obtained 72 frozen tissues of primary and metastatic STLMS (cohort 1) for liquid chromatography-mass spectrometry (LC-MS) analysis (Figure 1A). For independent cohort 2, we obtained FFPE blocks of 219 STLMS cases and constructed tissue microarray blocks (Figure 1A). Patient demographics are summarized in Table S1. LC-MS analysis of global proteomes quantified 10,249 proteins in total and 7,325 proteins in at least 30% of samples (Table S2A). For phosphoproteomics, the analysis quantified 6,542 phosphorylated proteins in total and 4,584 in at least 30% of samples (Figure S1A, Table S2B). Batch correction removed TMT multiplexing effects from the data (Figure S1B). Using non-negative matrix factorization (NMF) on filtered global proteome and phosphoproteome data, we identified three distinct proteome-based subtypes (P1, P2, and P3) of STLMS (Figures 1BC, S1CD). Survival analyses of patients who underwent complete surgical resection showed significant differences in overall survival (OS) and recurrence-free survival (RFS) among subtypes, with P2 showing the poorest survival. We next performed transcriptome-based subtype inference in the TCGA STLMS dataset as an orthogonal external comparison, in which inferred P2/P3-like tumors showed shorter recurrence-free survival and overlapped with TCGA iCluster1 (Fig. S1EF). Analysis of clinical attributes (Figure 1D) did not show significant differences in tumor size, age, tumor status (primary lesion or metastatic lesion), gender, TNM stage, or prior treatment between proteomic subtypes. However, metastatic samples were enriched in P2/P3 subtypes compared to P1 (p=0.0308). Primary tumor site distribution varied significantly among subtypes (p=0.01735), with trunk samples predominantly in P1 and pelvis samples in P2. FNCLCC Grade 1 tumors were only seen in P1, while FNCLCC Grade 3 tumors were only seen in P3.

**Figure 1.**
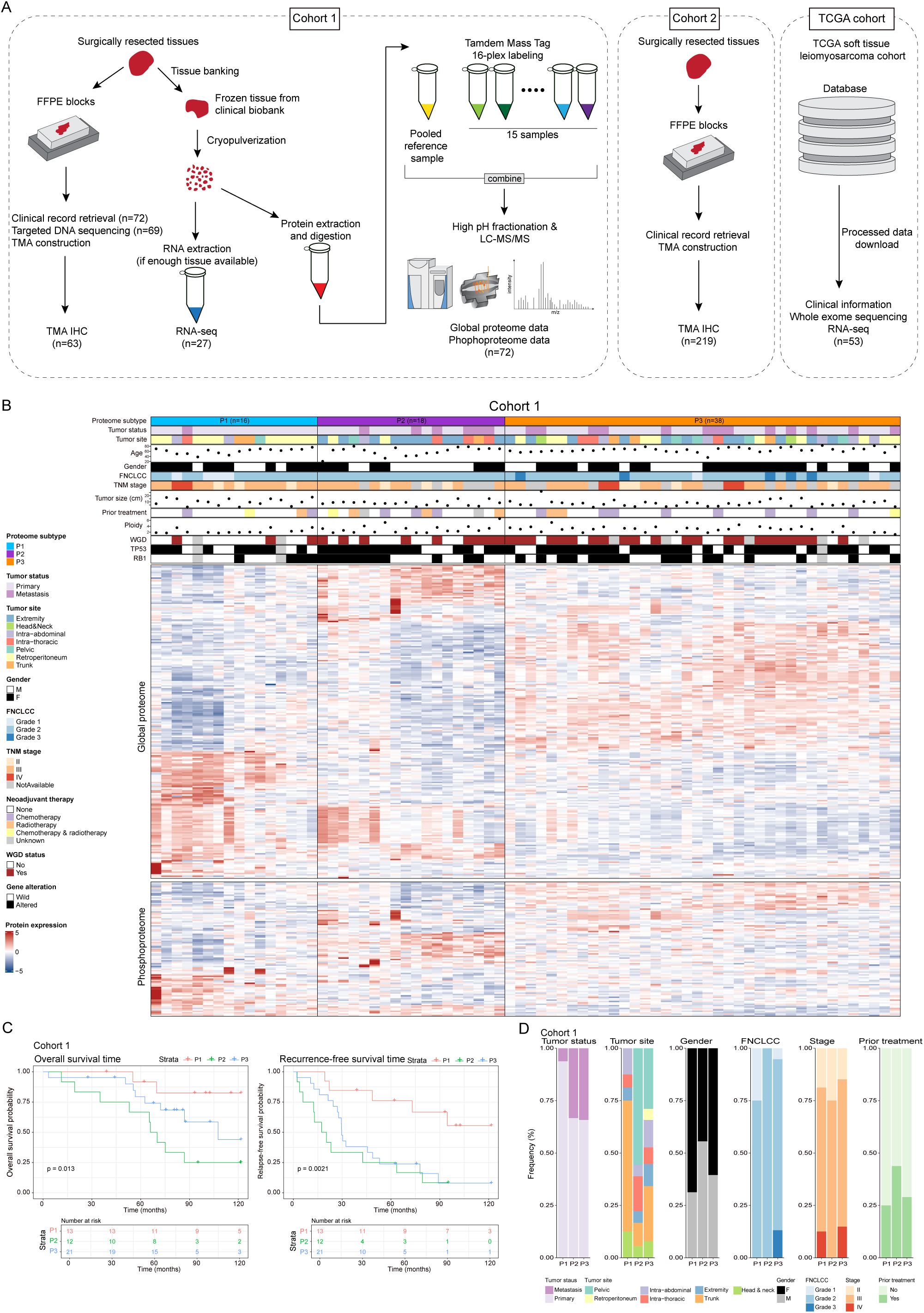
Integrative study design and clinicopathological features of proteome subtypes. (A) Summary of the workflow and metrics of this study. (B) Summary plot of cohort 1 including clinicopathological information and proteome subtypes. In the heatmap, proteins and phosphoproteins that show significant differences (FDR <0.05) between proteome subtypes are shown. (C) Kaplan-Meier curves of overall survival and recurrence-free survival between proteome subtypes. (D) Differences of clinicopathological factors between proteome subtypes.

### Differences of proteogenomic landscape and transcription regulatory networks between proteome-based subtypes

We evaluated genomic alterations in 69 samples by MSK-IMPACT targeted sequencing. Notably, P2/3 subtypes showed higher ploidy, whole genome duplication, and chromosomal instability versus P1 (Figure 2AB, findings supported by analysis of the TCGA STLMS cohort (Figure S2AB). Genomically, *TP53* (68%) and *RB1* (41%) were most frequently altered, followed by *NCOR1* (35%), *MAP2K4* (33%), and *FLCN* (28%) (Figure 2C). The GENIE STLMS cohort (v15.0, n=774)^21^ showed similar frequencies. On the other hand, P2/P3 showed frequent genomic alterations in *NCOR1*, *MAP2K4*, *FLCN*, and *IGF1R*. P2 showed commonly alterations in oncogenic drivers like *IGF1R* or *PDGFRA* and SWI/SNIF complex genes, associated with poor outcomes^22–25^. Somatic copy number alteration (SCNA) analysis revealed more arm-level amplifications in P2/P3 than in P1, while P3 showed higher deletion frequencies (Figure 2DE). Each subtype showed unique focal amplification/deletion patterns, with P1 showing fewer focal peaks (Figure 2F). The 17p12 peak, containing *NCOR1* and *MAP2K4*, was shared between P2 and P3. *NCOR1* and *MAP2K4* alterations occurred frequently together (p=1.03E-14, Fisher’s exact test). We analyzed gene overlap in recurrent focal peaks, finding 1,818, 1,125, and 3,556 involved genes, respectively (Figure 2G). Notably, Gene ontology analyses showed P1 focal deletion peaks enriched in copper/zinc ion processes, P2 focal peaks in extracellular matrix processes, and P3 focal peaks in T-cell receptor signaling (Figure S2C). KEGG analysis showed P2/P3 deletion peaks enriched in immune-related pathways, while only P3 showed deletions involving in PI3K-Akt oncogenic signaling (Figure S2D).

**Figure 2.**
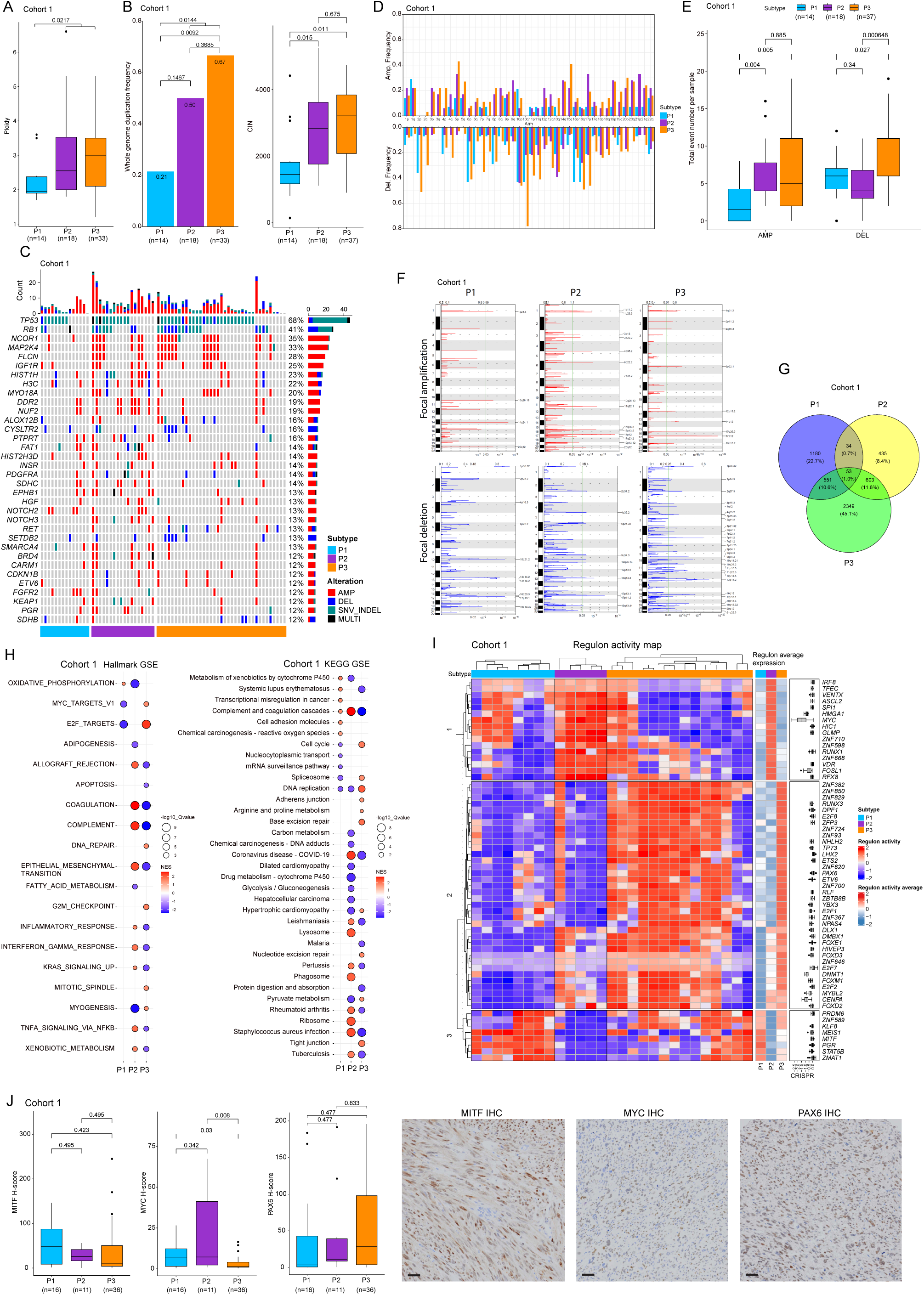
Landscape of genomic alterations and regulatory activity across proteome subtypes. (A) A boxplot showing ploidy between subtypes. P2/P3 show significantly higher ploidy than P1. (B) Bar chart of whole genome duplication (WGD) frequency in each subtype on the left and boxplot of chromosomal instability on the right. P2/P3 show significantly higher frequency of WGD and a higher CIN score than P1. (C) Oncoprint of integrated genomic alterations including SNV, Indel, and CNA. Genes with 8 events across cohort 1 (>12% samples) are shown. The bar chart on the right side of the oncoprint shows the alteration frequency across cohort 1. The bar chart on the top shows the number of genomic events of each sample. (D) Bar plot showing arm-level amplifications and deletions. In general, P1 shows lower frequency of arm-level events for each chromosome. P3 shows the highest frequency of arm-level deletions for most chromosomes. (E) Concordant with arm-level event frequencies for each chromosome, total event numbers of amplifications are significantly higher in P2/P3 than in P1. For arm-level deletions, P3 shows the highest event numbers compared with P1 or P2. (F) Focal amplification/deletion patterns in each subtype are shown. P1 has a relatively lower number of focal peak events. The 17p12 peak (NCOR1 and MAP2K4 gene loci) is shared between the P2 and P3 subtypes. (G) Venn diagram showing overlap of genes in in the recurrent focal peaks of each subtype. (H) Balloon plot showing pathway enrichment analyses of Hallmark and KEGG terms between subtypes. Only statistically significant results are shown. (I) Regulatory activity profile of STLMS by subtype status based on transcriptome data. Computed regulon activity scores are shown as a heatmap. Regulons with p<1E-8 from the Master Regulator Analysis process are shown. The boxplot on the right side of the heatmap shows DepMap CRISPER gene effect scores for 36 sarcoma cell lines. (J) Boxplots of MYC, PAX6, MITF IHC scores by subtypes are shown. In addition, representative IHC images are shown. Each transcription factor shows an expression profile of subtypes that is concordant with regulon network analyses of transcriptome data. Bars, 50 μm.

Pairwise differential expression analyses between subtypes (P1 vs. P2/P3, P2 vs. P1/P3, P3 vs. P1/P2) revealed 2,277 dysregulated proteins in P1 vs. others (207 upregulated >2-fold, 211 downregulated >2-fold), 2,503 in P2 vs. others (282 upregulated >2-fold, 286 downregulated >2-fold), and 2,652 in P3 vs. others (106 upregulated >2-fold, 377 downregulated >2-fold) with q-values <0.05 (Figure S2E) (Table S3ABC). KEGG and Hallmark analyses identified cancer-related pathways in each subtype (Figure 2H). P1 showed low proliferation and enriched metabolic processes. P2 showed enrichment of epithelial-mesenchymal transition (EMT), NF-κB pathway, inflammatory responses, and KRAS signaling, with downregulated oxidative phosphorylation. P3 exhibited cell cycle activation, DNA repair, low EMT, and higher muscle differentiation. Transcriptome regulatory network analysis identified 60 dysregulated regulons across proteome subtypes (Figure 2I). The regulons formed 3 clusters. Cluster 1 showed high activity in the P2 subtype with poor outcome (Figure 1B), with MYC enrichment confirmed by IHC (Figure 2J). Cluster 2 showed high activity in P3, with enriched cell cycle-related transcription factors. DNMT1 enrichment in P3 suggests methylation differences. PAX6, a tumor progression factor^26–28^, was upregulated in P3 and confirmed by IHC (Figure 2J). Cluster 3 showed high activity in P1 and some P3 samples. MITF, regulating immune response^29^, was enriched in P1 (Figure 2J), which is also supported by TCGA STLMS dataset (Figures 2J, S2F).

### Impact of genomic alterations on proteomic subtypes and cell cycle-related components

We studied *cis* effects of copy number alteration (CNA) on mRNA and protein. Analysis found 320 and 365 *cis* correlations of CNA-mRNA and CNA-protein respectively (FDR <0.1) (Figure 3A), with 131 genes overlapping (Figure 3B). GOBP over-representation analysis of cis regulated genes that showed positive CNA-protein abundance correlations revealed significant enrichment of cell cycle-related terms (Figure 3C). Most cell cycle components showed positive correlation with the cell cycle score at CNA, mRNA, and (phosho)protein levels (Figure 3D). The P3 subtype showed highest cell cycle activity, Ki67 immunolabeling (Figure 3E), and AURKA protein abundance (Figure 3F), an emerging cancer treatment target^30^. E2F targets were enriched in P2/P3 (Figure 3G), with CDK1/2 kinase activity significantly higher than in P1 (Figure 3H). Independent TCGA STLMS data provided orthogonal support for these findings (Figure S3A-C). DepMap CRISPR data showed that CDK1 and AURKA perturbation reduced cell viability in STLMS sarcoma cell lines, therefore supporting these molecules as candidate subtype-associated dependencies that warrant functional testing in LMS-specific models (Figure 3I). Metastatic STLMS samples showed increased cell cycle score, proliferation rate, and CDK2 kinase activity relative to primary site STLMS (Figure S3DE), with a trend for elevated AURKA protein expression (Figure S3F). IHC scores in the independent TMA-based cohort 2 supported these findings (Figure S3G). Combined cohort analyses of recurrence-free survival showed poor outcomes associated with increased expression of cell cycle-related components (Figure S3H), supporting further evaluation of AURKA- and CDK-associated programs in aggressive STLMS.

**Figure 3.**
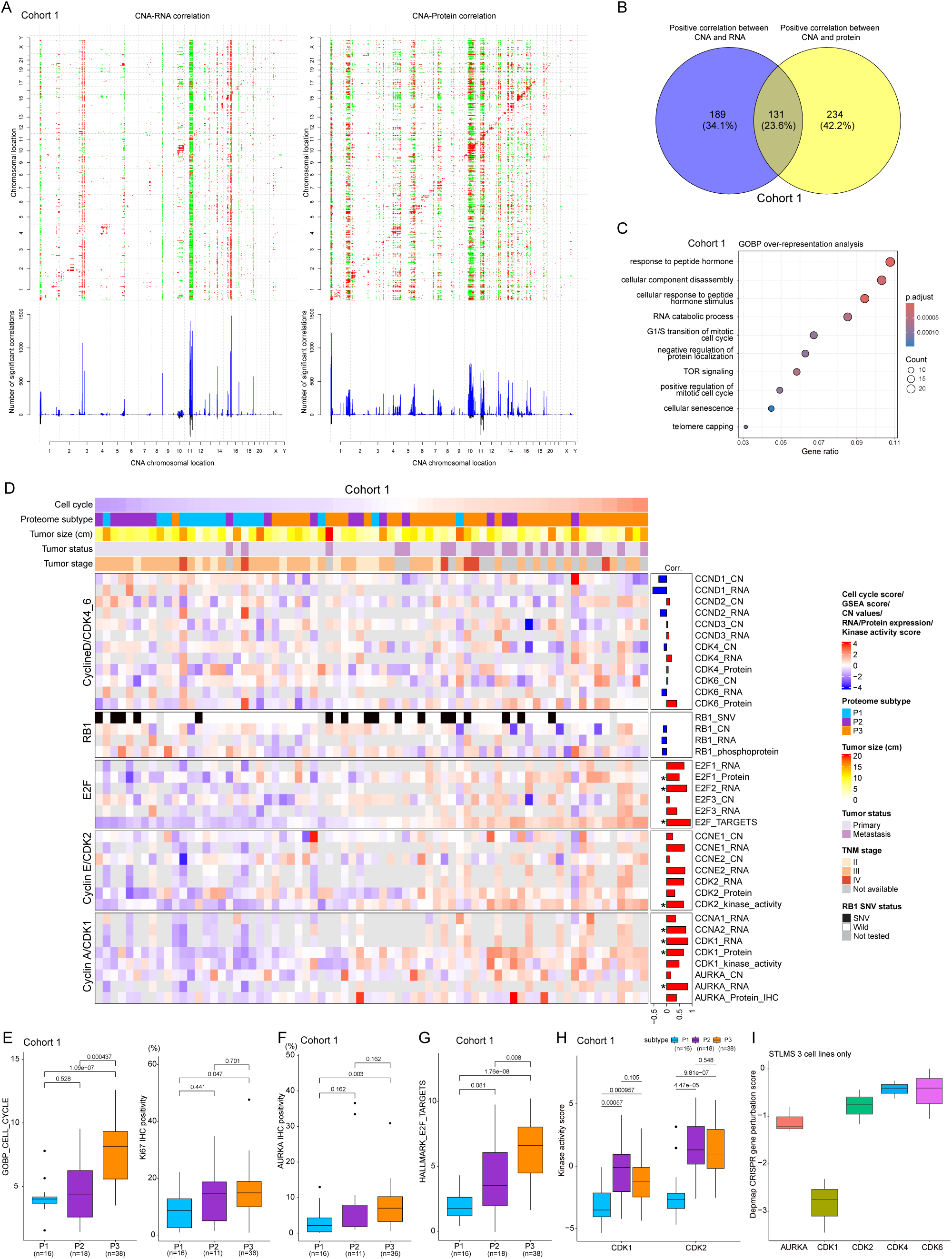
Impact of genomic alterations on the proteome. (A) The correlations between CNA and mRNA (left) and protein expression (right) levels (y axes) reveal both cis and trans effects. The presence of significant positive (red) and negative (green) Spearman’s correlations (FDR <0.1) between CNA and mRNA or protein is highlighted. The numbers of significant cis and trans events corresponding to the indicated genomic loci specific to mRNAs or proteins are represented by blue bars, while the overlap between CNA-mRNA and CNA-protein events is depicted by black bars. (B) Venn diagram showing cis correlation overlap between CNA-mRNA and CNA-protein. (C) GOBP over-representation analysis of cis correlation between CNA-protein is shown. Cell cycle-related terms are significantly enriched. (D) Genomic effect of cell cycle-related genes on mRNA and protein. Columns represent each sample and are ordered by cell cycle score. The bar chart on the right side of heatmap shows Spearman’s correlation coefficients between targets and cell cycle score. “E2F targets” is based on ssGSEA of the Hallmark E2F term. Kinase activity scores (CDK1,2 kinase activities) are from PTM-GSE analysis. p_RB1 denotes phosphorylated protein of RB1. * denotes statistical significance (p<0.05). (E) Boxplots showing proteome subtype differences of GOBP cell cycle scores and Ki67-positive tumor frequency. (F) Boxplot showing AURKA protein expression (IHC score) by subtypes. (G) Boxplot of E2F scores from ssGSEA by subtypes. (H) Boxplot showing kinase activity scores of CDK1/2 by subtypes. (I) Boxplot of Depmap CRISPR gene perturbation scores targeting CDKs and AURKA using 3 leiomyosarcoma cell lines. Score below -1 represents significant effect on cell survival when the target gene is inhibited.

### Phosphoproteomic differences across proteomic subtypes and receptor tyrosine kinase signaling

We investigated phosphoproteomic differences across STLMS subtypes focusing on receptor tyrosine kinase molecules (RTK-RAS pathways). Among 69 DNA-sequenced samples, 56 samples (81.2%) had at least one genomic alteration in RTK-RAS pathway genes. *MAP2K4* and *IGF1R* were the most frequent events. Though alteration frequencies were higher in P2/P3 vs. P1, differences were not statistically significant (Figure 4A). Genomic alteration frequencies of *PDGFRA* (overall frequency,14%), *FGFR2* (12%), *MET* (9%), *TSC2* (7%), *ERBB3* (6%), *FGFR4* (6%), *INPP4B* (6%) differed significantly among proteomic subtypes. All these genes except *FGFR2* showed enrichment in P2, while *FGFR2* was enriched in P1. Differential phosphoproteomic analysis across subtypes showed enrichment of Ras protein signal transduction in P2 and cell cycle-related terms in P3 (Figure 4B), consistent with the RTK–RAS genomic alterations observed in P2 and the cell cycle-related features shown in Figures 2 and 3. We next inferred kinase activity using phosphopeptide expression data and the RoKAI app^31^ to generate kinase-substrate networks (Figure 4CD). CSNK1A1 and PDK family kinases (PDK1, PDK2, PDK3, PDK4) were enriched in P1. AURKA activity was downregulated in P1, consistent with AURKA IHC assessment (Figure 3F). CDKs (CDK1, 2, 5, 6), AURKB, mTOR, and MAPK1 (ERK2) were enriched in P2. CDKs (CDK2, 7), AURKB, ROCK1, and BRAF were enriched in P3. AKT1/3 also showed enrichment in P3 versus P1/P2, although this difference did not reach statistically significance. Overall, P1 showed a distinct kinase activity profile from P2/P3, whereas P2 and P3 shared kinase activities such as AURKB or CDK2. This difference may reflect distinct genomic alteration patterns across subtypes (Figure 4A). PTM-GSEA showed similar subtype-associated differences in kinase activity (Figure S4A). AURKB kinase activity scores were higher in P2/P3 than P1 (Figure 4E). Consistently, AURKB IHC positivity was higher in P2/P3 than in P1 (Figure 4F), and AURKB mRNA expression in the TCGA cohort showed the same pattern (Figure S4B). According to PhosphoSitePlus database (https://www.phosphosite.org/homeAction), T232 is a well-characterized activating phosphorylation site of AURKB kinase. Motif analysis centered on AURKB-T232 identified candidate upstream kinases associated with this phosphosite pattern (Figure 4G). Most candidate upstream kinases were more active in P2/P3, whereas several AURKB-associated kinases showed lower activity in P1.

**Figure 4.**
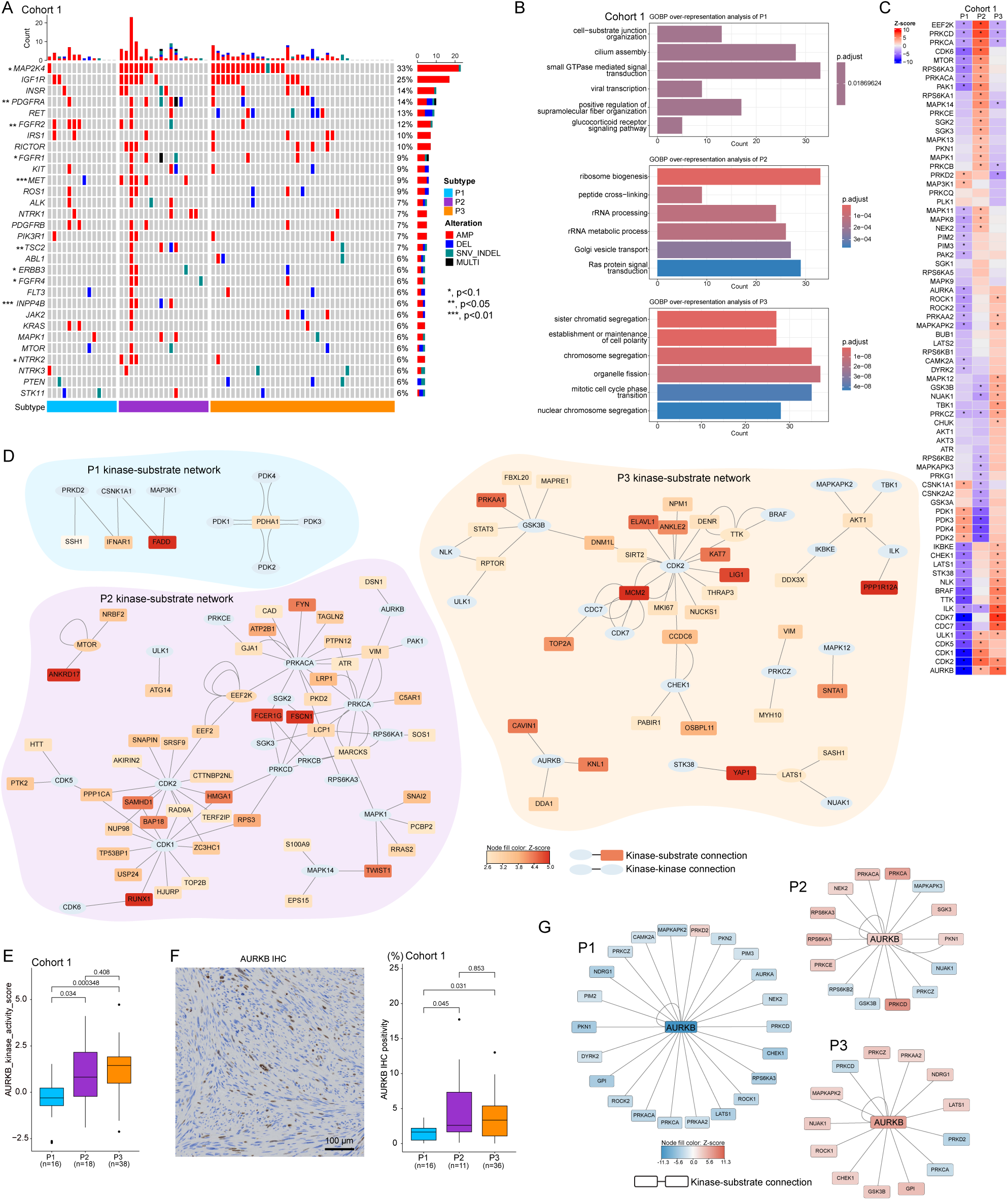
Phosphoproteomic differences between subtypes. (A) Oncoprint showing RTK-RAS pathways with 4 or more events. Amplification of MAP2K4 and IFG1R are the top 2 most frequent alterations. * denotes statistically significant differences (Fisher’s exact test) in gene alteration frequency between subtypes. (B) Over-representation analyses of GOBP terms. Phosphoproteins which are positively upregulated in one subtype compared to other subtypes with FDR<0.05 are used for GOBP term enrichment analyses. (C) Kinase activity heatmap by subtypes. Z-scores of inferred kinase activities are plotted as a heatmap. * denotes statistical significance. Kinase names are shown on the left side of the heatmap. (D) Kinase-substrate networks of the top 20 positively enriched kinases in each subtype are shown. Note: all positively enriched kinases of P1 are shown due to its small number. Genes shown as ellipses are kinases. Genes shown as rectangles are kinase substrates Colors of kinase substrates denote the phosphorylation Z-score of the target’s phosphorylation site. If multiple phosphorylation sites exist in a substrate, phosphorylation averages are shown based on color scale. (E) Boxplot of PTM-GSEA AURKB activity score shows enrichment in P2/P3 compared to P1. (F) Representative AURKB IHC image and boxplot of AURKB IHC score by subtype. Bar,100 µm. (G) Networks of AURKB and inferred regulatory kinases are shown. Motif analysis of AURKB-T232 identifies 194 kinases. Among them, kinases with significant enrichment (FDR<0.05) in RoKAI app analysis between subtypes were selected to be shown here. Colors of kinases denote Z-scores of kinase activity.

### NCOR1 expression is associated with progenitor-like and differentiation-related features in STLMS

Epigenetic regulators influence gene expression through multiple chromatin-related mechanisms, including histone modification, DNA methylation, transcription factor binding, and chromatin accessibility^32^. Given the frequent NCOR1 amplification observed in our cohort (Figure 2C), we next examined genomic alterations in epigenetic regulator genes in cohort 1 (Figure 5A). *NCOR1* showed the highest alteration frequency (35%), followed by *SMARCA4* (13%), *BRD4* (12%), and *CARM1* (12%). Alteration frequencies of *BRD4*, *NSD1*, and *SMARCA4* differed significantly among proteomic subtypes. *NCOR1* amplification was observed in more than 30% of P2/P3 tumors, compared with 14.2% of P1 tumors (Figure 5B). *NCOR1* copy number correlated with NCOR1 protein abundance, with P3 showing a trend toward the highest expression. Similarly, P2/P3 tumors in the TCGA STLMS cohort showed higher frequencies of *NCOR1* amplification, and *NCOR1* mRNA expression was highest in P3 (Figure S5A, B). Because *NCOR1* was one of the most frequently altered epigenetic regulators and was enriched in the poor-outcome P3 subtype, we further examined molecular features associated with high versus low NCOR1 protein expression. Tumor samples were divided into 2 groups by median NCOR1 LC-MS based protein level, followed by differential expression analysis and gene set enrichment analysis (GSEA). GSEA showed enrichment of muscle organ development-related terms in the NCOR1-high group, whereas immune-related and epithelial process-related terms were relatively depleted (Figure 5E). Smooth muscle differentiation markers also showed higher expression in the NCOR1-high group, although some differences did not reach statistical significance (Figure 5F), and similar patterns were observed in the TCGA cohort (Figure S5C). In addition, the NCOR1-high group showed higher stemness scores (Figures 5G and S5D), together with increased expression of PROM1 (CD133) and ALDH7A1 at the protein level (Figure 5H). In the TCGA cohort, the NCOR1-high group also showed higher mRNA expression of *POU5F1* (*OCT3/4*), *PROM1* (*CD133*), and *ALCAM* (Figure S5E). CD133 immunohistochemistry in TMA cohort 2 showed a similar trend, with higher CD133 expression in the NCOR1 IHC-high group than in the NCOR1 IHC-low group. YAP1/TEAD-related signals were also higher in the NCOR1-high group (Figure 5I, J), and similar enrichment of YAP1/TAZ (encoded by *WWTR1*)/TEAD-related features was observed in the TCGA dataset (Figure S5F). Together, these findings indicate that high NCOR1 expression is associated with progenitor-like and differentiation-related molecular features in STLMS; however, functional studies will be required to determine whether NCOR1 directly regulates these programs.

**Figure 5.**
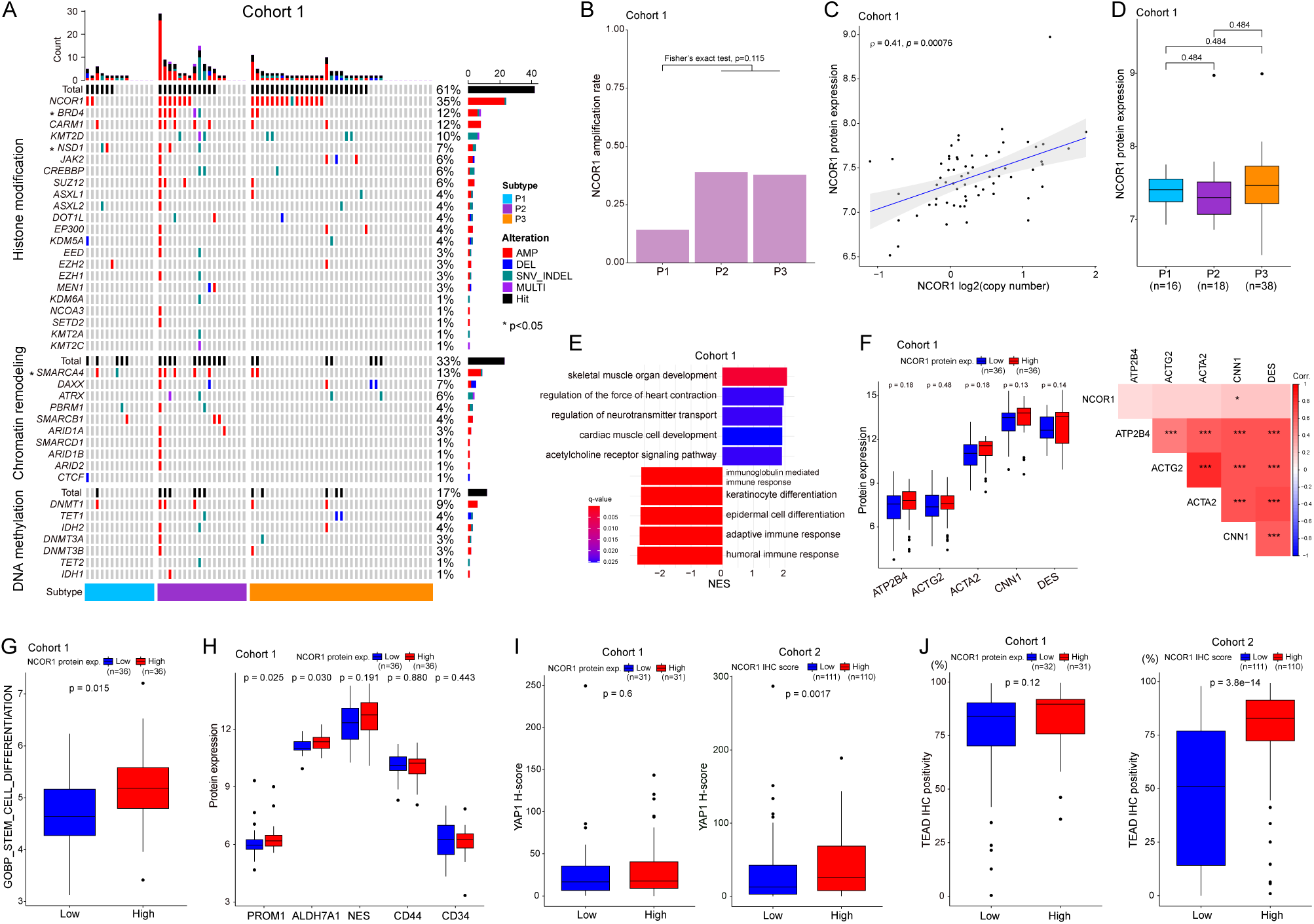
NCOR1 expression corelates with muscular differentiation, stemness, and immune status. (A) Oncoprint of epigenetic regulators in cohort 1. NCOR1 gene is the top altered epigenetic regulator. * denotes statistical difference of gene alteration frequency between subtypes. (B) NCOR1 amplification shows higher frequency in P2/P3 compared to P1 in cohort 1 (but not statistically significant). (C) NCOR1 copy number positively correlates with its protein level (Spearman’s correlation coefficient) in cohort 1. (D) Boxplot of NCOR1 protein expression by subtypes in cohort 1. (E) Top 5 positively and negatively enriched GOBP GSEA results are shown. A ranked gene list based on fold change values between NCOR1-low and -high groups in cohort 1 was used for GSEA. (F) Boxplot of smooth muscle protein markers between NCOR1-low and NCOR1-high. (G) Boxplot of GOBP_STEM_CELL_DIFFERENTIATION ssGSEA scores. In cohort 1, the NCOR1-high group shows significantly higher scores compared to the NCOR1-low group. (H) Boxplot of stemness markers between NCOR1-low/high groups. PROM1 (CD133) and ALDH7A1 in the NCOR1-high group show significantly higher protein expression than the NCOR1-low group in cohort 1. (I) Boxplots of YAP1 IHC H-scores between NCOR1-low and -high groups in cohorts 1 and 2. (J) Boxplots of TEAD IHC H-scores between NCOR1-low and -high groups in cohorts 1 and 2.

### Enrichment of homologous recombination deficiency and nonhomologous end joining-related features in poor-outcome subtypes

We observed increased chromosomal instability (CIN) in P2/P3 (Figures 2B and S2B), together with whole-genome duplication and enrichment of DNA repair-related programs in P3 (Figure 2H). Because homologous recombination repair (HRR) is a major pathway for the repair of DNA double-strand breaks (DSBs)^33^, and its impairment has been associated with genomic instability and adverse clinical behavior in multiple tumor types^34–36^, we next investigated HRR-related alterations in STLMS. Using a set of 64 HRR pathway genes defined from the TCGA sarcoma cohort^37^, we identified alterations in 38 genes in cohort 1. Integration of allele-specific copy number and variant data showed frequent loss of heterozygosity (LOH) affecting HRR pathway genes (Figure 6A), with HRR-related events more commonly observed in P2/P3 than in P1 (Figure 6B). The homologous recombination deficiency (HRD) score was highest in P2 (Figure 6C) and positively correlated with CIN (Figure 6D). Similar patterns were observed in the TCGA dataset (Figure S6A–D).

**Figure 6.**
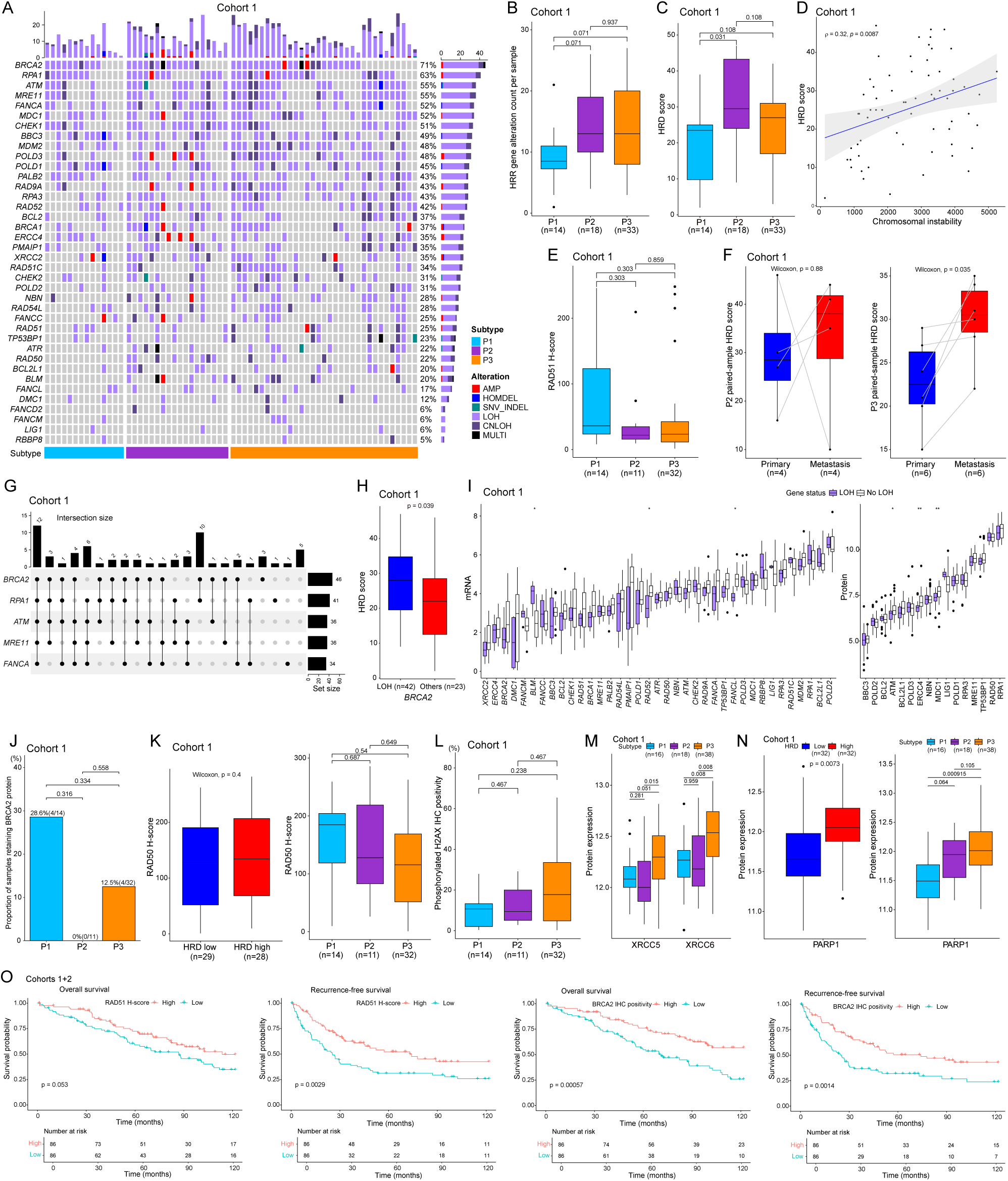
Homologous recombination repair pathway. (A) Oncoprint of HRR pathway components showing frequent LOH events in cohort 1. (B) Boxplot of genomic alteration counts in the HRR pathway per sample shows high occurrence in P2/P3, but not statistically significant. (C) The HRD score is higher in P2/P3 compared to P1. (D) The HRD score positively correlates with CIN. (E) The RAD51 IHC scores are lower in P2/P3 compared to P1, which is concordant with the HRD scores of these subtypes. (F) Boxplot of HRD score comparison between paired primary and metastatic samples of P2 and P3. Metastatic samples of P3 show a significant increase. Although overall not statistically significant, 3 out of 4 samples of P2 show HRD score increases. (G) UpSet plot of thetop 5 altered genes. *BRCA2* is the top altered gene with multiple different cooccurrence partners. (H) Boxplot of HRD scores between *BRCA2* LOH and non-LOH groups showing significant increase of HRD scores in the LOH group. (I) Boxplots of mRNA and protein expression between LOH and non-LOH groups for each HRR gene. Among 37 HRR genes (*FANCD2* was excluded due to low LOH event number), 16 proteins were quantified in cohort 1 and are shown here. (J) Bar plot showing frequencies of BRCA2-retaining samples in each subtype. P2/P3 subtypes, especially P2, show decrease of BRCA2 retention compared to P1. (K) The RAD50 IHC score does not change significantly between HRD-low and HRD-high groups, but it shows a downward trend in P2/P3 compared to P1. (L) Boxplot of phosphorylated H2AX IHC scores by subtypes. (M) Boxplot of XRCC5/6 protein expression by subtypes. P3 shows significantly higher expression of XRCC5/6 compared to P1 or P2. (N) Boxplot of PARP1 protein expression between HRD-low and HRD-high groups by subtypes. (O) Kaplan-Meier analysis of overall and recurrence-free survival in combined cohorts 1 and 2, stratified by median values of RAD51 H-score and BRCA2 tumor cell positivity.

To further assess HRR-related features, we examined RAD51 protein abundance by IHC, as RAD51 is a key recombinase in homologous recombination^38^. RAD51 staining showed a trend toward lower expression in P2/P3 than in P1 (Figure 6E), and this pattern was also supported by TCGA STLMS RPPA data (Figure S6E). In paired samples from the same patients, metastatic lesions in P3 showed higher HRD scores than matched primary lesions (Figure 6F), consistent with the association between genomic instability and metastatic progression reported in prior studies^35,36^.

We next examined genomic alteration patterns in the five most frequently altered HRR genes in STLMS, namely BRCA2, RPA1, ATM, MRE11, and FANCA, in cohort 1. The most frequent pattern involved concurrent alteration of all five genes (Figure 6G). Among single-gene and two-gene events, BRCA2 alterations were the most common (total events = 17). Tumors with BRCA2 LOH showed higher HRD scores than BRCA2-intact tumors (Figure 6H). More broadly, assessment of the remaining 37 HRR genes showed that LOH-positive tumors tended to have lower mRNA and protein expression of the affected genes (Figure 6I), a pattern that was also supported by the TCGA STLMS mRNA dataset (Figure S6F). TCGA STLMS RPPA data further showed reduced BRCA2 protein levels in HRD-high tumors and in BRCA2-LOH tumors (Figure S6G). In our cohort, BRCA2 loss by IHC was frequent in P2 (Figure 6J), which was also the subtype with the highest HRD score (Figure 6C).

We also evaluated RAD50 protein abundance by IHC because RAD50 is a core component of the MRN complex involved in early DSB recognition. Although RAD50 expression did not differ significantly according to HRD status alone in cohort 1, P2/P3 showed lower RAD50 expression than P1 (Figure 6K), and a similar pattern was observed in the TCGA dataset (Figure S6H). Assessment of phosphorylated H2AX, a marker of DNA damage signaling, showed increased positivity in P2/P3 relative to P1 (Figure 6L), consistent with altered DSB-associated signaling across subtypes.

We next examined non-homologous end joining (NHEJ), an alternative pathway for DSB repair. Specifically, we assessed classical NHEJ components KU70/KU80 (encoded by XRCC6/XRCC5) and the alternative NHEJ-associated factor PARP1^39^. XRCC5/6 expression was increased in P3 relative to P1, whereas P2 did not differ significantly from P1 (Figure 6M). PARP1 expression was higher in the HRD-high group and in P3 (Figure 6N), with P2 also showing a near-significant increase, consistent with increased expression of selected NHEJ-related components, particularly in P3. Similar findings were observed in the TCGA dataset (Figure S6I). Finally, using RAD51 and BRCA2 IHC data from cohorts 1 and 2, we found that lower expression of both proteins was significantly associated with poorer patient outcome (Figure 6O).

### Tumor immune microenvironment profiling identifies subtype-associated immune states in STLMS

To characterize immune microenvironment differences across STLMS, we performed integrative proteogenomic analyses together with IHC-based assessment of immune cell infiltration. Immune microenvironment status was evaluated using the ESTIMATE ImmuneScore^40^, which correlated with CD3+ T-cell counts by IHC (Figure 7A). The ImmuneScore also correlated with antigen-presenting machinery (APM) components, including MHC class I pathway members and IRF1 (Figure 7A). In addition, APM-related molecules showed concordance across genomic copy number, mRNA, and protein expression (Figure 7A), suggesting that genomic alterations may contribute to immune microenvironment differences in STLMS. The P3 subtype showed a significantly lower ImmuneScore (Figure 7B), a finding that was also supported by the TCGA dataset (Figure S7A, B). Paired-sample analyses further showed that metastatic tumors had lower ImmuneScores (Figure 7C) and reduced expression of immune-related processes (Figure S7C).

**Figure 7.**
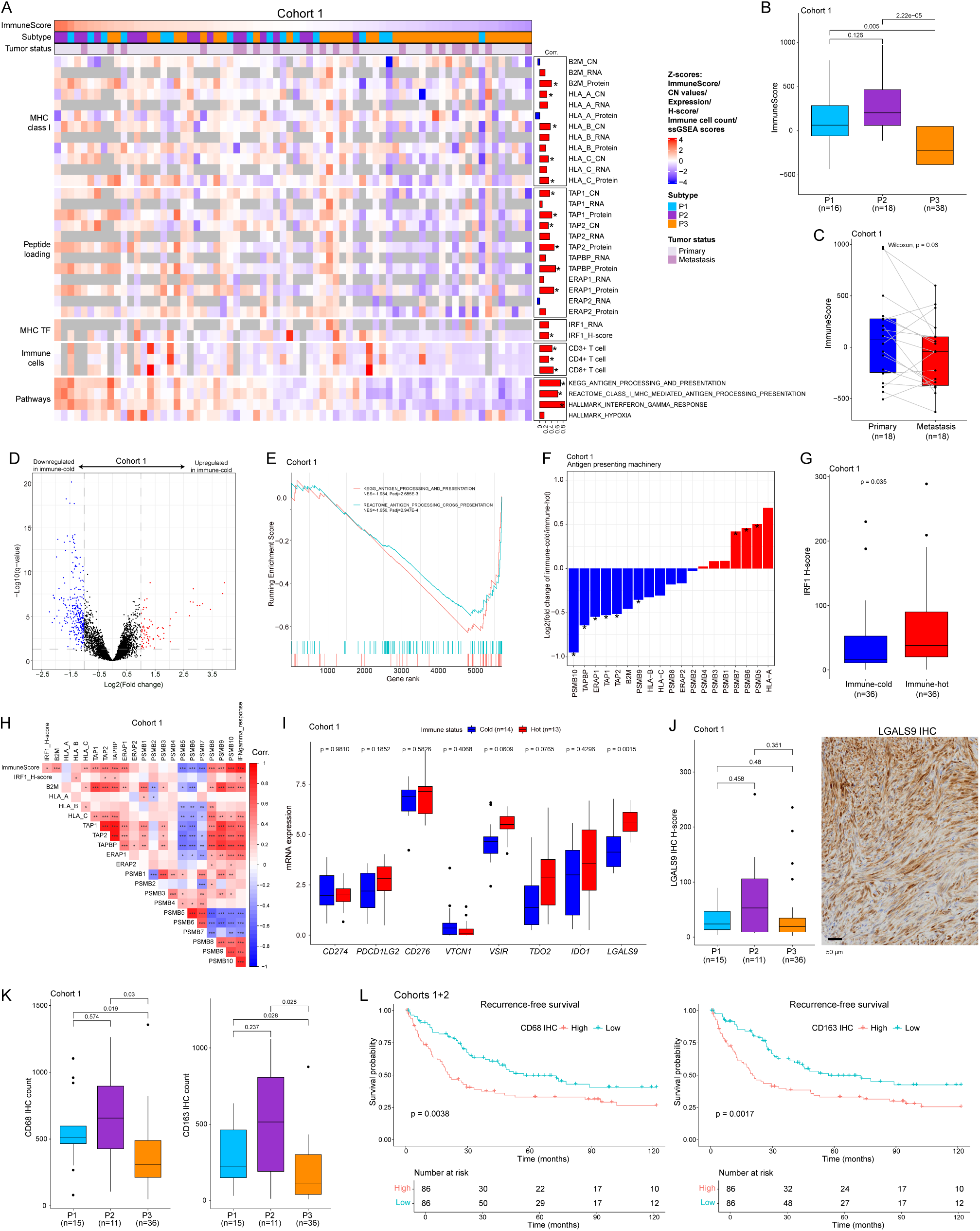
Tumor microenvironment profiling shows subtype differences and immune evasion characteristics in both immune-cold and immune-hot tumors. (A) A heatmap of immune signatures including copy number status, mRNA, and protein level. In general, the ImmuneScore positively correlates with CD3 count, antigen presenting molecules, and immune-related pathways. Bars on the right side of the heatmap denote Spearman’s correlation coefficients between ImmuneScores and each gene/signature. (B) Boxplot showing the ImmuneScore by proteome subtypes for cohort 1. (C) Boxplot showing the ImmuneScore between paired primary and metastatic tumors. (D) Volcano plot showing differential protein expression results between immune-cold and immune-hot. Proteins with statistical significance (q<0.05) are shown in red (log2(fold change)>1) and blue (log2(fold change) < -1). The horizontal dashed line denotes q=0.05. (E) GSE plot of antigen presenting pathway-related KEGG and REACTOME terms, showing antigen presenting processes are significantly downregulated in immune-cold tumors compared to immune-hot tumors. (F) A bar plot of APM protein expression ratios between immune-cold and immune-hot in cohort 1. * denotes statistical significance in the differential expressional analysis shown in (D). (G) Boxplot of IRF1 protein expression between immune-hot and immune-cold tumors. (H) A correlation plot (colored by Spearman correlation coefficient) including the ImmuneScore, IRF1, and antigen processing machinery protein expression. *, p<0.05; **, p<0.01; ***, p<0.001. IFNgamma_response scores are ssGSEA scores of the “Hallmark_Interferon_Gamma_Reponse” term. (I) Boxplot of immune-suppressive molecules between immune-cold and immune-hot tumors showing only TIM3 is significantly upregulated in immune-hot tumors vs. immune-cold tumors. (J) Boxplot of LGALS9 protein expression by subtypes (left side) and representative IHC image (right side). P2 shows a trend for higher expression of LGALS9 compared to P1 or P3. Bar, 50 µm. (K) Boxplots of CD68-positive or CD163 -sitive cell counts by subtypes. (L) Kaplan-Meier analyses of recurrence-free survival in combined cohorts 1 and 2, stratified by median values of CD68-positive or CD163-positive cell counts.

Comparison of immune-cold and immune-hot tumors identified 356 differentially expressed proteins, including 75 upregulated and 281 downregulated in immune-cold tumors (Figure 7D and Table S4). GSEA showed downregulation of antigen-presentation-related pathways in immune-cold tumors (Figure 7E), which was also supported by the TCGA STLMS dataset (Figure S7D). Metastatic tumors showed similar downregulation of antigen-processing pathways (Figure S7E). ERAP1, TAP1/2, MHC class I complex molecules (HLA-B/C and B2M), and proteasome components were downregulated in immune-cold tumors (Figure 7F). IRF1, a key transcriptional regulator of MHC class I and APM molecules^41^, also showed lower expression in immune-cold tumors (Figure 7G). We observed positive correlations between IRF1 and MHC class I molecules (HLA-B/C and B2M), although these correlations did not reach statistical significance. IRF1 also correlated positively with proteins involved in antigen processing, including TAP1, TAP2, TAPBP, ERAP1, PSMB8, and PSMB9, whereas the catalytic proteasome subunits PSMB5 and PSMB7 showed negative correlations (Figure 7H). Similar findings were observed in the TCGA transcriptomic dataset (Figure S7F). In addition, IRF1 expression correlated with the IFNγ-JAK-STAT pathway, consistent with its known upstream regulation^42^. By contrast, hypoxia score did not show a significant correlation with the STLMS ImmuneScore (Figure 7A).

We next examined the expression of immunoregulatory molecules in immune-hot tumors, which may attenuate antitumor immune responses. Transcriptomic analysis of cohort 1 showed that LGALS9 was upregulated in immune-hot tumors (Figure 7I). Consistently, the TCGA STLMS dataset showed increased expression of several immunoregulatory molecules, including LGALS9 (Figure S7G). Because LGALS9 was identified in both cohorts, we further evaluated its protein expression by IHC in cohorts 1 and 2. IHC showed LGALS9 expression in tumor cells, with a trend toward higher expression in the P2 subtype, which was relatively immune-hot (Figure 7J). Notably, LGALS9 protein expression was not detected by IHC in benign smooth muscle tissues, including vascular and intestinal smooth muscle. A similar pattern was observed in the TCGA STLMS cohort (Figure S7H).

To further characterize the immune microenvironment of P2, we assessed tumor-associated macrophages by IHC using CD68 as a pan-macrophage marker and CD163 as an M2-like macrophage marker. CD163-positive macrophages were enriched in the P2 subtype (Figure 7K). In survival analyses, higher CD68-positive or CD163-positive cell counts were associated with shorter recurrence-free survival (Figure 7L). Together, these findings indicate that the P2 subtype is associated with an immune-hot yet potentially suppressive microenvironment characterized by higher LGALS9 expression and increased M2-like macrophage infiltration. However, these data do not establish that LGALS9 directly mediates macrophage recruitment or immune evasion, which requires future in-depth investigation.

### Development and validation of an IHC-based classifier for proteome-defined STLMS subtypes

To enable practical subtype assignment, we developed a TMA-based IHC classifier for the three proteome-defined STLMS subtypes P1, P2, and P3 (Figure 8A). The candidate pool included 23 IHC markers, comprising 18 markers assessed in this study and 5 additional subtype-informative markers identified through subtype-specific differential expression analyses comparing each subtype against the other two subtypes combined. Using cohort 1 as the training set, we systematically tested all 3-, 4-, 5-, and 6-marker combinations by multinomial logistic regression with 10-fold cross-validation and selected the optimal model on the basis of subtype-discriminative performance. This approach identified a 6-marker classifier consisting of CD133, MYC, CDK2, AURKA, AURKB, and HLA-ABC (Figure 8A). The final model used a multinomial logistic regression framework with a predefined probability threshold for subtype assignment; cases not meeting the threshold were designated unclassified. The 6 markers showed subtype-associated expression patterns in both cohort 1 and cohort 2, supporting reproducibility across independent TMA cohorts (Figure 8B). When the locked classifier was applied to cohort 2, the inferred proteomic subtypes showed significant differences again in recurrence-free survival (Figure 8C, log-rank p = 0.039). P1 was associated with the most favorable outcome, whereas P2 and P3 showed less favorable recurrence-free survival.

**Figure 8.**
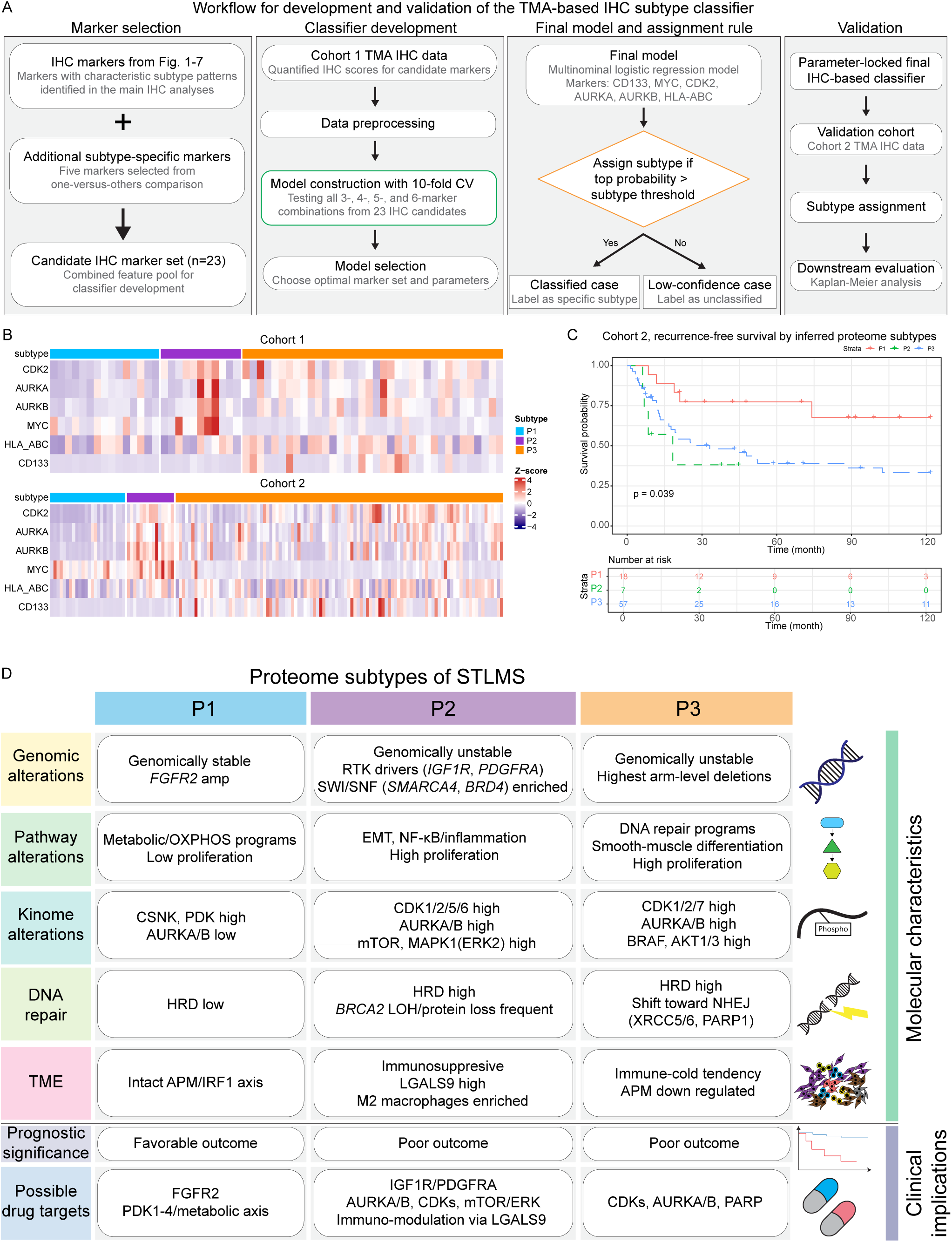
IHC-based classifier development and integrated summary of proteome-defined STLMS subtypes. (A) Workflow for development of an IHC-based classifier for proteome-defined STLMS subtype prediction. Twenty-three candidate IHC markers were evaluated, including 18 markers assessed in this study and 5 additional subtype-informative markers identified through subtype-specific differential expression analyses. All possible 3-, 4-, 5-, and 6-marker combinations were tested in the training cohort using multinomial logistic regression with 10-fold cross-validation. The final selected classifier included CD133, MYC, CDK2, AURKA, AURKB, and HLA-ABC. (B) Heatmap of the 6 selected IHC markers in cohort 1 and cohort 2. (C) Recurrence-free survival according to inferred proteomic subtype in the independent validation cohort 2 classified by the locked IHC model. (D) A schematic overview of the proteome-defined STLMS subtypes summarizing major genomic alterations, pathway and kinome features, DNA repair states, tumor microenvironment characteristics, prognostic associations, and potential therapeutic vulnerabilities. APM, antigen-processing machinery; EMT, epithelial-mesenchymal transition; HRD, homologous recombination deficiency; LOH, loss of heterozygosity; NHEJ, nonhomologous end joining; TME, tumor microenvironment.

To summarize the major biological and clinical features of each subtype, we generated an integrative schematic of the proteome-defined STLMS subtypes (Figure 8D). This overview highlights subtype-associated genomic alterations, pathway and kinome features, DNA repair states, tumor microenvironment characteristics, prognostic associations, and potential therapeutic vulnerabilities. Collectively, these findings support the clinical feasibility of IHC-based subtype assignment and provide a concise framework for biological interpretation of the three proteome-defined STLMS subtypes.

## Discussion

In this integrative proteogenomic analysis of STLMS, we identified three proteome-defined subtypes (P1, P2, and P3) with distinct molecular and clinicopathologic features (Figure 8). These subtypes were associated with differences in patient outcome, genomic alterations, signaling pathway activity, and tumor immune microenvironment, providing a framework for understanding biological heterogeneity in STLMS.

Unsupervised clustering identified P2 as the subtype with the poorest outcome, showing the shortest overall survival (OS) and recurrence-free survival (RFS). Importantly, survival analyses were restricted to primary tumors with complete (R0) resection, supporting that the adverse outcomes observed in P2/P3 are consistent with intrinsically aggressive tumor biology that is evident before metastasis. In the overall cohort, both P2 and P3 were enriched for metastatic lesions and higher-grade tumors, further supporting their association with clinically aggressive disease biology. External comparison using the TCGA STLMS dataset provided orthogonal support for this subtype framework. Although the TCGA study identified two major subtypes (iClusters 1 and 2)^12^, it lacked comprehensive phosphoproteomic profiling. Our integrated analyses identified three proteome-defined subtypes, providing an additional layer of molecular characterization of STLMS heterogeneity.

Genomic analyses revealed differences in genomic instability across subtypes. High ploidy and whole-genome duplication in P2 and P3 were consistent with a more genomically unstable phenotype associated with aggressive clinical behavior^43^. Increased chromosomal instability (CIN) in these subtypes may contribute to tumor heterogeneity and therapy resistance^44^. Alterations in oncogenic drivers such as IGF1R and PDGFRA were more frequent in P2/P3, while recurrent alterations in chromatin-regulatory genes, including SMARCA4, BRD4, NCOR1, and MAP2K4, pointed to subtype-associated differences in chromatin regulation and signaling programs. These findings support biologically distinct routes of tumor progression, although functional studies are needed to define their causal roles.

Copy number alterations correlated with mRNA and protein abundance, particularly for genes related to cell-cycle regulation (Figure 3). P3 showed the strongest cell-cycle activity together with elevated AURKA expression, nominating CDK- and Aurora-related programs as candidate dependencies in aggressive STLMS subtypes. DepMap CRISPR-based perturbation data support the biological relevance of some of these molecules^45,46^, although disease-specific validation in LMS models remains necessary.

The three subtypes also showed distinct pathway and kinome features. P1 was characterized by lower proliferation and metabolic programs. P2 showed enrichment of EMT, NF-κB signaling, immune-related programs, and RTK-RAS pathway features, whereas P3 was characterized by cell-cycle activation and DNA repair-related programs (Figures 2, 3, 6, and 7). Phosphoproteomic analyses identified subtype-associated kinase activity patterns consistent with these biological states. P1 showed relative enrichment of CSNK1A1 and PDK family kinases with lower AURKA/B activity. P2 was characterized by RTK-RAS/ERK-mTOR and cell-cycle-related kinase signals, including CDK1/2/5/6, AURKB, mTOR, and MAPK1 (ERK2). P3 showed CDK2/7, AURKB, ROCK1, and BRAF activity together with relatively increased AKT1/3 kinase activity. Across subtypes, AURKB kinase activity was higher in P2/P3 than in P1 and this pattern was supported by IHC (Figure 4E, F). Together, these kinome analyses nominate subtype-associated signaling dependencies for future functional testing.

Transcriptional regulatory network analysis identified distinct transcription factor programs across subtypes (Figure 2I). P2 showed higher activity of regulators including MYC, consistent with its aggressive phenotype. P3 showed enrichment of cell-cycle-associated regulators, including E2F family members, DNMT1, and PAX6. These observations provide a framework for future studies of subtype-specific regulatory circuitry in STLMS.

High NCOR1 expression was associated with increased smooth muscle marker expression together with higher expression of progenitor-like markers, including PROM1 (CD133) and ALDH7A1 (Figure 5). Increased YAP1/TAZ-TEAD-related signals in NCOR1-high tumors further support the possibility that differentiation-associated and progenitor-like programs coexist in a lineage-related tumor state. However, these observations are associative and do not establish a direct causal role for NCOR1 in stemness maintenance or smooth muscle differentiation^47^.

Our data also showed that HRD-related features were enriched in both P2 and P3, together with increased expression of selected NHEJ-related components (Figure 6)^48^. Elevated HRD scores were associated with increased chromosomal instability, and loss of heterozygosity in HRR genes, particularly BRCA2, was associated with reduced HRR-related expression. Increased phosphorylated H2AX positivity in P2/P3 was consistent with altered DNA damage signaling across subtypes (Figure 6L). PARP1 expression was also increased in HRD-high tumors and in P3, supporting the possibility that DNA damage response-targeted strategies may merit future evaluation in selected STLMS subsets. These implications remain hypothesis-generating until validated experimentally^49,50^.

Immune profiling showed that P3 and metastatic tumors had lower ImmuneScores together with downregulation of antigen-presenting machinery components, supporting an immune-cold phenotype in these contexts. By contrast, immune-hot tumors showed increased expression of immunoregulatory molecules, including LGALS9, together with enrichment of M2-like macrophages. These findings suggest that STLMS contains biologically distinct immune states, including an immune-hot yet potentially suppressive microenvironment in P2. However, our data do not establish that LGALS9 directly drives macrophage recruitment or immune evasion.

To advance the clinical translation of the proteomic subtype framework, we developed a tissue microarray (TMA)-based immunohistochemistry (IHC) classifier utilizing a limited marker panel and applied the established model to an independent validation cohort. The inferred subtypes demonstrated significant differences in recurrence-free survival, thereby supporting the feasibility of approximating proteome-defined STLMS subtypes through a pathology-based surrogate assay. Although further refinement and external validation are necessary prior to clinical application, this approach offers a practical bridge between proteogenomic subtype discovery and clinically deployable patient stratification.

In conclusion, this study offers a comprehensive proteogenomic characterization of STLMS, integrating phosphoproteomic analysis of subtype-associated kinase programs. By delineating proteome-based subtypes with distinct genomic, signaling, DNA repair, and immune characteristics, this research serves as a valuable resource for future biological and translational investigations of STLMS. The findings highlight CDK/Aurora-related signaling, HRD-associated features, and subtype-specific immune states as potential areas for further exploration. However, additional orthogonal validation, mechanistic studies, and further refinement and external validation of practical surrogate assays are required before these findings can be applied to clinical stratification or treatment selection in routine practice.

## Materials and Methods

### Clinical specimens and pathological data

We examined tissues from two cohorts of STLMS patients. Cohort 1 consists of 72 primary and metastatic STLMS tissues from 54 patients, including 18 primary-metastasis-paired samples. Cohort 2 consists of 219 surgically resected STLMS samples. The study was approved by the Institutional Review Board (MSKCC #16-1683, BIDMC #2023P000933). Clinical data, such as patient demographics, treatment history, recurrence status, clinical targeted MSK-IMPACT sequencing results, or histologic grade were retrieved from medical records in an anonymized fashion. Tumor content ratios for all samples were verified by board-certified pathologists. Clinicopathologic features and prognosis information, including gender, age, pathological TNM stage, anatomic site, are summarized in Table S1.

### Targeted cancer gene sequencing

MSK-IMPACT targeted sequencing was conducted as previously outlined^51^. In summary, genomic DNA was extracted from both tumor and adjacent normal tissues using the DNeasy Tissue kit and the EZ1 Advanced XL system (Qiagen). The extracted DNA was fragmented with the Covaris E200 device. Custom DNA probes were crafted for targeted sequencing of all exons and selected introns across 505 genes. These probes were synthesized using the NimbleGen SeqCap EZ library custom oligo system and were biotinylated. Sequencing libraries were prepared following the KAPA HTP protocol (Kapa Biosystems) and the Biomek FX system (Beckman Coulter), and sequencing was performed on the Illumina HiSeq 2500 to achieve high, uniform coverage (>500x median coverage). Demultiplexed fastq files underwent trimming with Trim Galore v0.2.5mod (https://github.com/FelixKrueger/TrimGalore) to remove adapters and short reads. These files were then aligned to the human reference genome (GRCh37) using BWA-MEM v0.7.5a (with arguments -M -t 6)^52^, and Picard Tools v2.9 (https://github.com/broadinstitute/picard) was used. AddOrReplaceReadGroups was applied to annotate read groups, and MarkDuplicates was used to identify PCR duplicates. Genomic regions were pinpointed using FindCoveredIntervals from the GATK Tool Kit v3.3-0^53^and underwent indel realignment with Assembly Based ReAligner (ABRA) v2.12^54^. GATK BaseRecalibrator was employed to identify systematic errors in base quality scores. Variant calling was executed in paired tumor/normal mode using MuTect v1.1.4^55^for single nucleotide variants (SNV). Pindel v0.2.5a7^56^was utilized for small insertions and deletions (indels). Vardict v1.5.1^57^served as the variant caller to report both SNVs and indels. The vcfs generated by MuTect, Vardict, and Pindel were subsequently combined. Additionally, copy-number variants, including chromosomal instability (CIS) and whole-genome doubling (WGD), were identified using FACETS^58^. The resulting variants were annotated using vcf2maf v1.6.14 (https://github.com/mskcc/vcf2maf), which employs Ensembl’s Variant Effect Predictor v86. In conclusion, we successfully conducted MSK-IMPACT sequencing on 69 out of 72 samples in cohort 1 (LM_010P, LM_020P, LM_043P lacked sufficient DNA for sequencing). Amplifications and deep deletions were identified from the GISTIC result table named “allthresholded.bygenes.txt” (part of the GISTIC output used to determine the copy number status of each gene in each sample). Genes with a value of +2 were classified as amplifications, while those with a value of -2 were considered deep deletions. Sequencing results are detailed in Table S5.

### Allele-specific copynumber analysis

FACETS analyses were conducted to ascertain genome-wide allele-specific and absolute DNA copy numbers for all target sequencing samples except 4 samples (FACETS version 0.5.6)^58^. Allele-specific copy number data were utilized to estimate tumor purity and ploidy. Prior to further analyses, total copy number log ratios were adjusted for ploidy and purity. Tumors exhibiting whole-genome doubling (WGD) were identified as those in which more than 50% of the autosomal genome possessed a major copy number ≥2, where the major copy number is defined as the number of copies of the most prevalent allele present in the sample^59^. Loss of heterozygosity (LOH) was defined as the loss of one allele in cohort 1. For the TCGA STLMS cohort, allele-specific copy number data were obtained via the R TCGAbiolink package^60^, and LOH was defined using the same criteria.

### HRD score calculation

The scarHRD R package was utilized to calculate the HRD score of our cohort 1 samples^61^. This metric was derived from the simple addition of three types of genetic alterations: loss of heterozygosity (LOH), large-scale transitions (LST), and telomeric allelic imbalance (TAI) based on allele-specific copy number data. LOH refers to the elimination of genomic segments exceeding 15 Mb in size, excluding entire chromosomes. LST encompasses chromosomal breakages between adjacent regions of at least 10 Mb, with breaks occurring no more than 3 Mb apart. TAI quantifies the disparities in allele sequence contributions within the telomeric regions of chromosomes. For the TCGA cohort, we downloaded precalculated HRD score from the prior study^62^.

### Chromosomal instability inference

Chromosomal instability (CIN) scores of our cohort 1 and the TCGA STLMS data was computed on the allele-specific copy number segments by adding the number of gains (CNV >2), losses (CNV <2), and loss of heterozygosity per sample based on data from the prior study^37^.

### Recurrent somatic copy number alteration detection

Recurrent somatic copy number alterations (SCNAs) were identified using the Genomic Identification of Significant Targets in Cancer (GISTIC, version 2.0.23)^63^to ascertain which SCNA regions exhibited amplification or deletion at a frequency greater than expected by chance, with a q value threshold of 0.05. The analysis employed the following parameters: Amplification Threshold = 0.3, Deletion Threshold = 0.3, Cap Values = 1.5, Confidence Level = 0.99, Join Segment Size = 4, Arm Level Peel Off = 1, and Sample Normalization Method = mean. Default values were applied for all other parameters.

### RNA sequencing

For each sample, 20-30 mg of frozen tissue was homogenized in 1 mL of TRIzol Reagent (ThermoFisher catalog # 15596018), followed by phase separation induced with 200 µL of chloroform. RNA was extracted from 350 µL of the aqueous phase utilizing the miRNeasy Mini Kit (Qiagen catalog # 217004) on the QIAcube Connect (Qiagen) in accordance with the manufacturer’s protocol. Samples were eluted in 34 µL of RNase-free water. Subsequent to RiboGreen quantification and quality control via the Agilent BioAnalyzer, 500 ng of total RNA with RIN values ranging from 5.9 to 9.9 underwent polyA selection and TruSeq library preparation as per the instructions provided by Illumina (TruSeq Stranded mRNA LT Kit, catalog # RS-122-2102) with 8 cycles of PCR. Samples were barcoded and sequenced on a NovaSeq 6000 in a PE100 run, employing the NovaSeq 6000 S1 Reagent Kit (200 Cycles) (Illumina). Read quality was evaluated using FastQC. Trimmed raw reads were aligned to the human genome version hg19 using STAR (v2.7.5a)^64^with default parameters, and gene annotation was incorporated using Gencode v36, followed by the calculation of gene-level count values and TPM values via the RSEM tool (version 1.3.1)^65^.

### Tissue proteome extraction

Aliquots of 5 mg of frozen tissue were lysed using a lysis buffer composed of 8 M urea and 200 mM EPPS (4-(2-Hydroxyethyl)-1-piperazinepropanesulfonic acid), at pH 8.5, supplemented with protease inhibitor (complete mini EDTA-free, Roche) and phosphatase inhibitors (cocktails 2 and 3, Sigma). This was followed by homogenization using a Bead Ruptor. Subsequently, the tissue mixture underwent sonication through 12 cycles of 1-minute sonication at 120 W power (FB120, Fisher Scientific), with intermittent cooling. Post-centrifugation at 18,000 g for 10 minutes at 4 °C, the supernatant containing all soluble proteins was collected. Protein concentrations were quantified using BCA assays (Pierce). Pooled protein samples were prepared and stored, with equal amounts of all samples combined to assess MS run quality.

### Liquid chromatography-mass spectrometry (LC-MS) proteomic analysis

Protein samples were labeled utilizing the 16-plex TMT chemical labeling reagent (Thermo Fisher Scientific), followed by MultiNotch MS3 LC–MS analysis employing an Orbitrap Fusion MS^66^. In brief, 300 µg of proteins were reduced with 5 mM TCEP (tris(2-carboxyethyl) phosphine hydrochloride), alkylated with 10 mM IAA (iodoacetamide), and quenched with 10 mM DTT (dithiothreitol). The samples were subsequently diluted and precipitated using chloroform-methanol^67^. The resulting pellets were resuspended in 50 µL of 200 mM EPPS buffer and digested with Lys-C and trypsin at 37 °C overnight. Anhydrous acetonitrile was added to achieve a final volume of 30%. TMTPro (16-plex) reagents were introduced to the peptides at a 2.8:1 (TMT reagent-to-peptide) ratio and incubated for 1 hour at room temperature. A label ratio check was conducted to determine mixing ratios, labeling efficiency, and missed cleavages by pooling 1 µL from each sample. Based on these results, samples were mixed to achieve equal intensity across TMT channels. The samples were dried to remove acetonitrile, desalted using C18 solid-phase extraction (SPE) Sep-Pak (Waters), and vacuum-centrifuged. Five percent of the desalted samples were reserved for global proteome analysis, while 95% were utilized for phosphopeptide enrichment. Dried peptides intended for global proteomics were fractionated using the Pierce High pH Reversed-Phase Peptide Fractionation Kit (ThermoFisher catalog # 84868) into nine fractions per the modified manufacturer’s protocol. The fractions were vacuum-centrifuged and then reconstituted in 1% ACN/0.1% FA for LC–MS analysis. For optimal phosphopeptide enrichment, phosphopeptides were collected using the High-Select TiO2 Phosphopeptide Enrichment Kit (Thermo Fisher) and from the stored flow-through of Fe-NTA using the Pierce High-Select Fe-NTA Phosphopeptide Enrichment Kit (Thermo Fisher), and both were combined. Phosphopeptide samples were fractionated into nine fractions utilizing the Pierce High pH Reversed-Phase Peptide Fractionation Kit. The fractions underwent analysis via LC-MS/MS employing a nanoAQUITY UPLC (Waters) with a 50 cm (75 µm inner diameter) EASY-Spray column (PepMap RSLC, C18, 2 µm, 100 Å) maintained at 60 °C, in conjunction with an Orbitrap Fusion Lumos Tribrid Mass Spectrometer (Thermo Fisher). Peptide separation was conducted at a flow rate of 300 nL/min using a linear gradient of 1-35% acetonitrile (0.1% FA) in water (0.1% FA) over a duration of four hours, analyzed in SPS-MS3 (for global proteome) and neutral loss-triggered MS3 (phosphopeptide) modes. For global proteome analysis, MS1 scans were performed within a range of 375–1,500 m/z, with a resolution of 120 K, an AGC target of 4×1e5, and a maximum IT of 50 ms. MS2 scans were conducted on MS1 scans of charges 2-7 using 0.7 m/z isolation, collision-induced dissociation at 32%, turbo scan, and a maximum IT of 50 ms. MS3 scans employed specific precursor selection (SPS) of 10 isolation notches, an m/z range of 100–1,000, 45% CE, 50 K resolution, and an AGC target of 1e5. For phosphopeptide analysis, MS1 scans were conducted within a range of 380-1,400 m/z, with a resolution of 120 K, an AGC target of 4×1e5, and a maximum IT of 50 ms. MS2 scans were performed on MS1 scans of charges 2-7 using 0.7 m/z isolation, collision-induced dissociation at 35%, turbo scan, and a maximum IT of 50 ms. Neutral loss-triggered MS3 scans utilized a range of 100–1,000 m/z, 38% CE, 60 K resolution, an AGC target of 1e5, and a maximum IT of 118 ms. Targeted loss masses of 97.9763 and 79.9658 were employed.

### TMT data analysis

Raw data files were processed utilizing Proteome Discoverer (PD) version 2.4.1.15 (Thermo Scientific). In each of the TMT experiments, raw files from all fractions were consolidated and analyzed using the SEQUEST HT search engine with a Homo sapiens UniProt protein database, downloaded on 2019/12/13 (92,249 entries). Oxidation (M), phosphorylation (STY), and deamidation (NQ) were designated as variable modifications, whereas cysteine carbamidomethylation, TMT 16plex (K), and TMT 16plex (N-terminus) were specified as fixed modifications. The precursor and fragment mass tolerances were set at 10 ppm and 0.6 Da, respectively. A maximum of two missed trypsin cleavages were allowed. Searches employed a reversed sequence decoy strategy to control the peptide false discovery rate (FDR), with a 1% FDR established as the identification threshold. Phosphosite localization was determined using the PD IMP-ptmRS node. The raw abundance of protein, phosphoprotein, and phosphopeptide levels was normalized by sample loading normalization of each TMT channel after filtering out protein/peptide assignments with valid values for less than 30% of samples and K-Nearest Neighbors (KNN) imputation. Internal reference scaling based on pooled samples assigned for each TMT plex was conducted across the entire cohort to eliminate batch effects, followed by TMM normalization for median centralization between TMT batches^68,69^. Differential expression analyses were performed using edgeR^70^.

### Unsupervised clustering of proteomic data

We used non-negative matrix factorization (NMF)-based clustering of global protein and phosphoprotein expression data. In brief, we constructed a merged expression matrix of normalized global protein and phosphoprotein data and selected the top 500 (phospho)proteins with highest mean absolute deviations for unsupervised clustering to avoid noisy (phospho)proteins in the data. The resulting matrix was then subjected to NMF analysis leveraging the NMF R package^71^. To determine the optimal factorization rank *k* (number of clusters) for the matrix, we tested a range of clusters between *k* = 2-7. For each *k*, we factorized matrix *V* using 1,000 iterations with random initializations of W and H. To determine the optimal factorization rank, we calculated two metrics for each k: (1) cophenetic correlation coefficient (measuring how well the intrinsic structure of the data was recapitulated after clustering) and (2) silhouette score (indicating how well each sample lies within its cluster compared to other clusters^72^) (Figure S1CD). These metrics indicate the reproducibility of the clustering across 1,000 iterations. The cophenetic correlation coefficients for *k* = 2 and 3 were >0.95 and showed a large drop at *k* = 4. The silhouette score at *k* = 2, 3 showed relatively higher scores than that at *k* = 4. Based on these results and census heatmaps for each *k*, we chose *k* = 3 as the optimal clustering number with sufficient samples in each class for downstream analysis.

### Gene set enrichment analysis (GSEA)

Fold change values for gene expression between specified groups were calculated using edgeR. We employed clusterProfiler (v2.1.2)^73^with a ranked gene list ordered by fold change value. Enrichment tests were conducted for GOBP, Hallmark, KEGG, and Reactome gene sets obtained from the Molecular Signatures Database (MSigDB v7.4). For the functional characterization of each sample via single sample GSEA, we calculated normalized enrichment scores (NES) of cancer-relevant gene sets by projecting the matrix of normalized expression data onto GOBP, Hallmark, KEGG, and Reactome pathway gene sets. This was done using the ssGSEA implementation available at https://github.com/broadinstitute/ssGSEA2.0 with the following parameters: sample.norm.type = “log”, weight = 0.75, statistic = “area.under.RES”, output.score.type = “NES”, nperm = 1000, min.overlap = 3, correl.type = “z.score”. To elucidate gene set enrichment in each sample, we applied single sample GSEA on normalized expression data^74^and calculated the normalized enrichment score (NES) for MSigDB terms.

### Over-representation analysis (ORA)

ORA was performed to identify biological pathways and functional categories enriched in the gene set of interest by using clusterProfiler (v2.1.2)^73^. Enrichment was calculated using a hypergeometric test against the defined background, and p-values were adjusted for multiple comparisons using the Benjamini-Hochberg method.

### Posttranslational modification signature enrichment analysis (PTM-SEA)

To infer kinase activity based on phophosite-specific quantification data, we performed PTM-SEA (https://github.com/broadinstitute/ssGSEA2.0), an adaptation of ssGSEA, that analyzes site-specific signatures by scoring PTMsigDB bi-directional signature sets. PTMsigDB, sourced from over 2,500 publications, contains modification site-specific signatures linked to perturbations, kinase activities, and signaling pathways. Unlike other pathway databases, PTMsigDB annotates each PTM site with its directional change upon specific perturbations or signaling events, improving PTM-SEA’s scoring. PTMsigDB’s main source is the PhosphoSitePlus database^75^. The PTM-SEA requires two main inputs: a site-centric data matrix, denoted as m and formatted in GCT v1.3, along with the PTM signatures database. In matrix m, each row corresponds to a distinct phosphorylation site that is precisely localized to a particular amino acid residue, with the columns indicating the measured abundances across different samples. When multiple phosphorylation sites are identified on the same peptide, they are transformed into individual site-specific entities for each site.

### Kinase enrichment analysis between subtypes

To assess enrichment of activated kinases and phosphatases between groups based on functional information such as known protein-protein interactions or kinase-substrate annotations, we input phosphopeptide abundance change values (log2(fold change)) between proteome subtypes into the RoKAI application (v0.8.3)^31^ and obtained kinase activity scores at proteome subtype level. Default parameters were used.

### Transcriptional regulatory network analysis

To deduce the transcriptional regulatory activity of STLMS based on proteome subtype, we utilized the RTN R package^76–78^, which examines the link between specific transcription factors (TFs) or gene regulators and all possible targets, as well as the enrichment status of a network using proteome data. In summary, we employed normalized transcriptome data (n=27) and Lambert’s human TF list^79^(1,612 TFs) to initially build a potential transcriptional regulatory network (TRN). We then eliminated non-significant associations through permutation analysis (permutation n=1000) and bootstrapping, followed by the ARACNe algorithm^80^ to preserve direct TF-target interactions, thereby removing redundant indirect interactions. The refined TRN dataset includes the regulator gene name, its target gene name, and adjusted p-values as a confidence measure. For enrichment analyses over a list of regulons in the refined TRN, we first conducted a Master Regulator Analysis (MRA)^81^over a list of regulons, with adjustments for multiple hypothesis testing. The MRA evaluates the overlap between each regulon and the significantly dysregulated genes between subtypes, providing adjusted p-values, which resulted in 600 confident regulons (adjusted p-values <0.05) for further analyses. We then determined these regulons’ enrichment scores between proteome subtypes using transcriptome expressional fold change values and the GSEA-2T (two-tailed GSEA analysis) function from the RTN R package. To identify similarities and differences in regulatory programs among samples in our cohort, we selected 60 statistically robust regulons (adjusted p-values <1E-8), calculated the activity score of each regulon in each sample, and visualized the data as a heatmap. Additionally, we incorporated CRISPR perturbation scores (sourced from the Cancer Dependency Map (DepMap) Project^82^) of 36 sarcoma cell lines and included this data in the heatmap.

### Transcriptome-based inference of proteome-defined subtypes in the TCGA STLMS cohort

To perform an external comparison of the proteome-defined subtypes identified in cohort 1, we inferred subtype labels in the TCGA STLMS transcriptome dataset using a classifier trained on the subset of cohort 1 samples with matched transcriptome data and known proteome subtype assignments. Specifically, among cohort 1 samples, 27 cases had both transcriptome data and proteome-defined subtype labels (P1, P2, and P3) derived from proteomic and phosphoproteomic analyses. These 27 samples were used as the reference set for classifier construction. To select informative features, we first performed differential expression analyses across the proteome-defined subtypes within this 27-sample transcriptome subset and selected genes that were differentially expressed between subtypes at FDR < 0.01 and absolute fold change > 2. This yielded 668 genes for model construction. Normalized expression data for these genes were then divided into training (n = 19) and test (n = 8) sets, and a random forest classifier was trained using the tidymodels R package^83^ with 10-fold cross-validation. The classifier achieved an F-measure of 1.0 in the held-out test set. We then applied the trained model to the TCGA STLMS transcriptome dataset (n = 53) to assign transcriptome-inferred labels corresponding to the proteome-defined subtypes. TCGA sample IDs and inferred subtype assignments are listed in Table S6.

### Assessment of immune status

ImmuneScores were derived from the normalized protein expression matrix of 72 samples utilizing the ESTIMATE (Estimation of STromal and Immune cells in MAlignant Tumor tissues using Expression data) package in R^40^. To evaluate immune cell infiltration, we conducted IHC with immune cell markers (CD3, CD4, CD8, CD68, CD163) and tallied the number of positive cells for each marker across 3 TMA cores per sample. Subsequently, we computed the average number of positive cells per core for each sample for further analysis.

### Tissue microarray (TMA) construction

TMAs were created from FFPE blocks of 63 STLMS tissues in cohort 1 and 219 STLMS tissues in cohort 2. Three 1-mm cores were extracted from each tissue paraffin block and placed into tissue array blocks using a TMA arrayer (3DHistech). Tumor and normal regions were chosen after a thorough examination of individual histologic slides and electronic image-based selection of target areas for coring.

### Immunohistochemistry (IHC)

Four-µm sections were cut from FFPE tissue blocks. Paraffin was removed with xylene, and antigens were retrieval by heat-mediated epitope retrieval. Tissue sections were stained using a Leica BOND-MAX IHC autostainer. The antibodies used were anti-CD3 (Leica Microsystems, NCL-L-CD3-565, 1/200 dilution), anti-CD4 (Sigma, 104R-15, 1/100 dilution), anti-CD8 (Dako, M7103, 1/1500 dilution), anti-CD68 (Dako, M0876, 1/200 dilution), anti-CD163 (Leica Microsystems, NCL-CD163, 1/250 dilution), anti-MITF (Abcam, ab303530, 1/100 dilution), anti-MYC (Abcam, ab32072, 1/200 dilution), anti-PAX6 (Abcam, ab195045, 1/100 dilution), anti-AURKA (Cell Signaling Technology, #91590, 1/100 dilution), anti-AURKB (Abcam, ab45145, 1/100 dilution), anti-CDK1 (Abcam, ab133327, 1/250 dilution), anti-CDK2 (Cell Signaling Technology, #18048, 1/200 dilution), anti-active YAP1 (Abcam, ab205270, 1/20,000, dilution), anti-TEAD (Abcam, ab97460, 1/400 dilution), anti-NCOR1 (Invitrogen, PA1-844A, 1/1000 dilution), anti-RAD51 (Abcam, ab133534, 1/100 dilution), anti-BRCA2 (Sigma, HPA026815, 1/50 dilution), anti-RAD50 (Cell Signaling Technology, #61915, 1/4000 dilution), anti-phosphorylated H2AX (Cell Signaling Technology, #9718, 1/200 dilution), anti-Ki67 (Biocare, PRM325, 1/100 dilution), anti-IRF1 (Abcam, ab243895, 1/100 dilution), anti-LGALS9 (Abcam, ab227046, 1/200 dilution). For MYC, MITF, CDK1, CDK2, NCOR1, YAP1, RAD50, RAD51, PAX6, IRF1, and LGALS9 IHC. The staining intensity of each tumor cell was rated on a scale from 0 to 3+, and the average was taken from three separate tissue cores for each case. To determine the total weighted IHC score (IHC H-score, theoretically ranging from 0 to 300) for a sample, the expression intensity of individual tumor regions (rated 0-3+) was multiplied by their respective contributions (0-100%) to the overall tumor area, and these values were then summed. Two pathologists independently evaluated all tissue samples. If there were any discrepancies, the cases were re-evaluated until a consensus score was achieved. For Ki67, AURKA/B, TEAD, BRCA2, and phosphorylated H2AX IHC, we assessed the fraction of positive tumor cells. BRCA2 loss was defined as less than 50% positivity.

### Development of an IHC-based classifier for LMS subtype prediction

To develop an IHC-based classifier for LMS subtype assignment, we used the discovery cohort with matched proteomic subtype labels as the training set. A total of 23 candidate IHC markers were evaluated, including 18 markers assessed in this study and 5 additional subtype-informative markers identified through differential expression analyses comparing each subtype with the other two subtypes combined. Only cases with complete IHC data were included in model development. All possible combinations of 3, 4, 5, and 6 markers were systematically evaluated using multinomial logistic regression models to predict the three LMS subtypes P1, P2, and P3. Model performance was assessed by 10-fold cross-validation, and candidate marker sets were ranked according to subtype-discriminative performance. To improve assignment confidence, a rejection rule was applied such that a subtype was assigned only when the highest predicted probability exceeded a predefined subtype-specific threshold; otherwise, the case was labeled unclassified. Subtype-specific probability thresholds and a margin threshold were optimized within the training cohort. Final model selection was based on 10-fold cross-validation metrics, including macro-F1 score, classification coverage, accuracy among classified cases, and the number of unclassified cases. The final selected model included 6 markers, CD133, MYC, CDK2, AURKA, AURKB, and HLA-ABC, with subtype-specific probability thresholds of 0.73 for P1, 0.65 for P2, and 0.57 for P3. In the training cohort, this model classified 43 of 62 cases, corresponding to 69.4% coverage, with 79.1% accuracy and a macro-F1 score of 0.613 among classified cases. The locked 6-marker classifier was then applied to the independent validation cohort for downstream analyses. The final classifier model, preprocessing files, and prediction script are available in a GitHub repository (https://github.com/atanakapth/lms-ihc-classifier).

### Survival time analysis

Prognostic associations of genes and proteins of interest with overall survival (OS) and recurrence-free survival (RFS) were assessed using Kaplan–Meier survival analysis. For each target gene or protein, patients were classified into groups based on the predefined expression cutoff or genomic status. The analysis was restricted to patients with primary lesions who underwent R0 surgical resection. Differences in OS and RFS between groups were compared using the log-rank test. Owing to the limited sample size, TNM stage-specific and tumor site-specific analyses were not performed. Statistical significance was defined as a two-sided *p* < 0.05.

### External datasets

The TCGA STLMS dataset (53 samples) including clinical information, transcriptome data (count data, TPM), and gene alterations with allele-specific copy number data (data for 1 sample was not available) was obtained via the TCGAbiolink R package (version 2.30.0)^60^. Protein expression data from reverse phase protein array (RPPA) experiments of the TCGA cohort was downloaded from MD Anderson Cancer Center database (https://tcpa.drbioright.org/rppa500/main.html).

## Data and code availability

Our mass spectrometry proteomics data have been deposited in Harvard Dataverse and is available under controlled access (DOI: 10.7910/DVN/BWOCM9). Raw and processed data of MSK-IMPACT genomic sequencing and transcriptome data were deposited to the EGA repository with the identifiers EGAS50000000594 and EGAS50000000595, respectively. Other data derived from this study will be shared upon reasonable request. We did not generate original code in this study. Any additional information required to reanalyze the data reported in this paper is available from the lead contact upon request.

## Abbreviations (there are a lot more missing)

STLMS: soft tissue leiomyosarcoma
IHC: immunohistochemistry
SNV: single nucleotide variant
FFPE: formalin fixed paraffin embedded
GSEA: gene set enrichment analysis
PTM-SEA: posttranslational modification signature enrichment analysis
CIN: chromosomal instability
CNA: copy number alteration
DSB: DNA double-strand break
EMT: epithelial–mesenchymal transition
FDR: false discovery rate
APM: antigen-presenting machinery.

## Acknowledgements

MHR expresses gratitude for the grants received from the NIH/NCI (R21 CA251992, R21 CA263262, and U01 CA263986), a Cycle for Survival Equinox Innovation Grant, an Investigator Grant from the Neuroendocrine Tumor Research Foundation (NETRF), and the support from the Farmer Family Foundation. Portions of the research were funded by the MSKCC NCI Cancer Center Support Grant (P30 CA008748). The funding organizations did not participate in the study’s design, nor in the data collection, analysis, or interpretation.

## Author Contributions

AT designed the study, carried out experiments, performed data analyses, and wrote a draft of the manuscript. MO performed sample processing and clinical data collection. YO carried out experiments and data analysis. RH and ZL supported mass spectrometric experiments and data analyses. NA provided expert advice on sarcoma pathology. DK consulted on the study and provided partial funding. JW and WW wrote and edited the manuscript. MR supervised the study, analyzed data, wrote and edited the manuscript, and provided funding.

## Declaration of generative AI and AI-assisted technologies in the manuscript preparation process

During the preparation of this work the authors used Paperpal in order to check English grammar. After using this tool/service, the authors reviewed and edited the content as needed and take full responsibility for the content of the published article.

## Supplemental information

Document S1. Figures S1-S7

Table S1. Clinicopathological characteristics of patients in cohort 1, related to Figures 1-7

Table S2. Proteome and phosphoproteome expression data of cohort 1, related to Figures 1-7

Table S3. Differential protein expression analysis between subtypes, related to Figure 2

Table S4. Differential protein expression analysis between immune-cold and immune-hot samples, related to Figure 7

Table S5. MSK-IMPACT summary with GISTIC 2.0 results of cohort 1, related to Figure 2

Table S6. Clinicopathological attributes and inferred proteome subtypes of the TCGA STLMS cohort, related to Figures S1-S7

---

**Figure S1.**
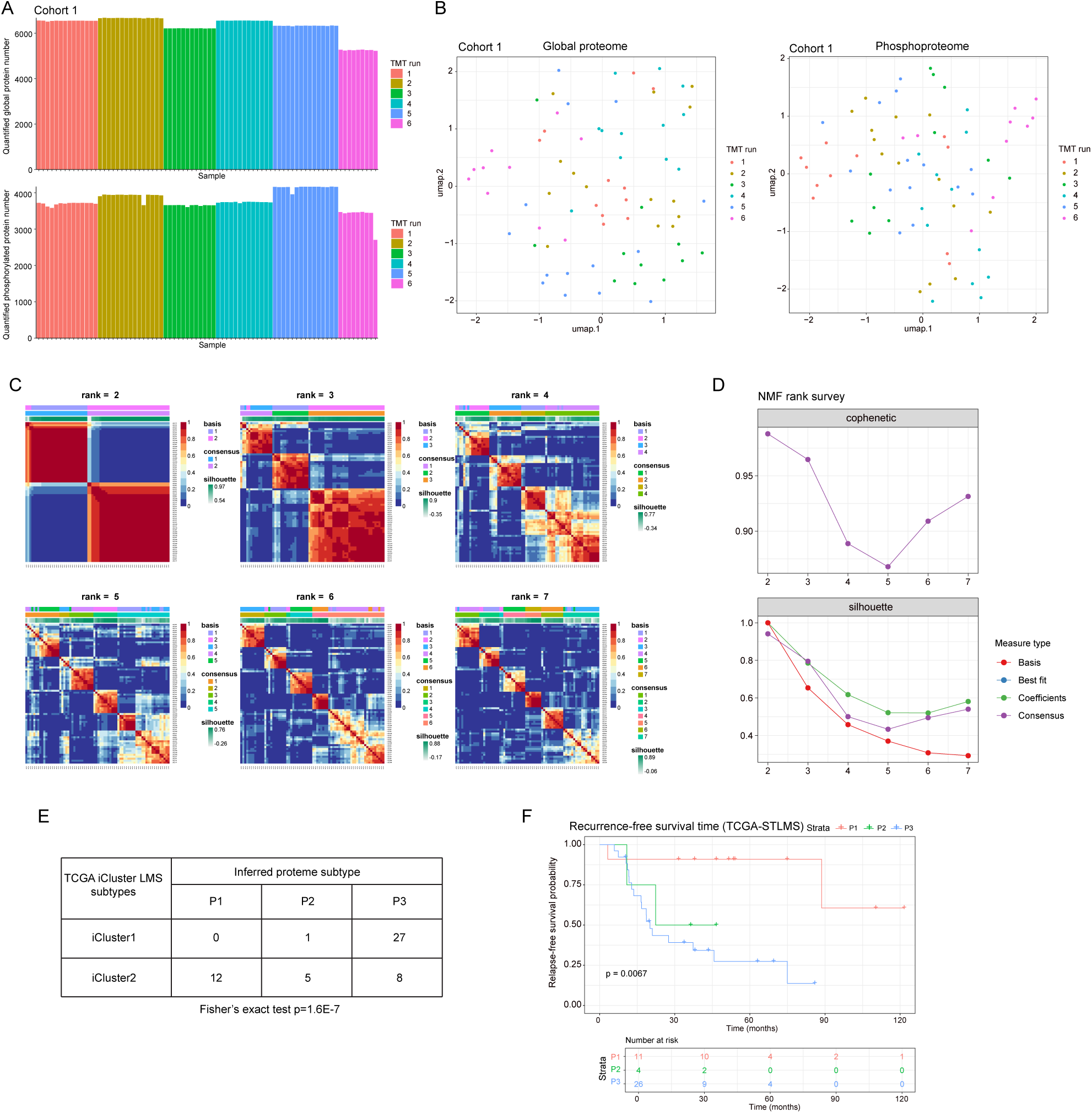
Evaluation of proteome data quality, batch effects, and robust subtype inference, related to Figure 1. (A) Bar plots showing the number of quantified proteins per sample for (top) global proteome and (bottom) phosphoproteome datasets. Samples are grouped and color-coded by TMT plex MS runs (1-6). The relatively consistent number of quantified proteins across samples and batches indicates robust and comparable proteome/phosphoproteome coverage across the TMT experiments. (B) UMAP plots of (left) global proteome and (right) phosphoproteome profiles, colored by TMT plex MS runs (n=6). Each point represents a sample. Samples are well intermixed across TMT batches, indicating no detectable batch effect in both global proteome and phosphoproteome datasets. (C) Consensus heatmaps for non-negative matrix factorization (NMF) clustering at ranks k=2-7, showing clustering stability across different cluster numbers. Each heatmap illustrates consensus values between sample pairs, with color bars indicating sample assignments in basis and consensus groupings. Silhouette width values are shown for each rank, reflecting clustering quality. (D) Summary plots of NMF rank survey metrics: cophenetic coefficient (top) and silhouette width (bottom) across ranks. Rank 3 shows a relatively high silhouette score and a drop in cophenetic coefficient, suggesting it as the optimal cluster number balancing stability and separation. (E) Association between proteome-defined subtypes (P1-P3) and TCGA iCluster LMS subtypes (iCluster1 and iCluster2) in the TCGA STLMS cohort. The distribution shows significant concordance between iCluster1 and subtype P3, and iCluster2 and subtypes P1/P2 (Fisher’s exact test, p=1.6×10E-7). (F) Kaplan-Meier analysis of recurrence-free survival in the TCGA STLMS cohort, stratified by proteome-based subtypes. Stratification by proteome-based subtypes shows significant survival differences between these subtypes. The P2/P3 subtypes show poorer recurrence-free survival compared to P1.

**Figure S2.**
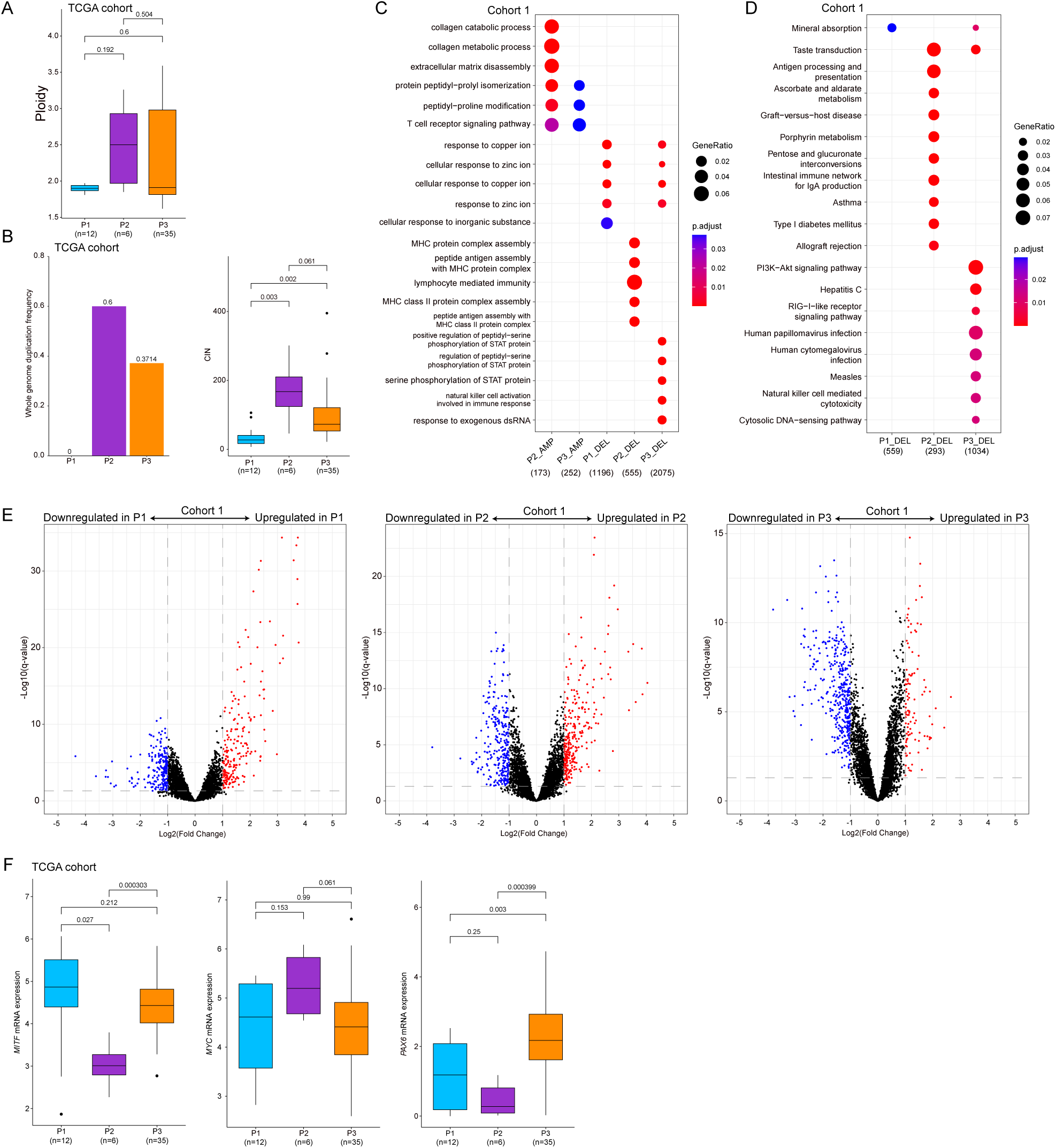
Genomic instability features and regulator profiles in the TCGA STLMS cohort and differential expression analyses of cohort 1, related to Figure 2. (A) Boxplot showing ploidy distributions by inferred proteome subtypes in the TCGA STLMS cohort (n=53). (B) Bar chart of whole genome duplication frequency by inferred proteome subtypes in the TCGA STLMS cohort (n=53) on the left and boxplot of CIN in the TCGA STLMS cohort (n=52) on the right. (C) Balloon plot showing GOBP over-representation analysis of genes involved in focal peak amplifications or deletions by subtype in cohort 1. (D) Balloon plot showing KEGG over-representation analysis of genes involved in focal peak amplifications or deletions by subtype in cohort 1. (E) Volcano plots showing differentially expressed proteins between subtypes in our cohort 1. (F) mRNA expression profile of representative regulons by subtypes in the TCGA STLMS dataset.

**Figure S3.**
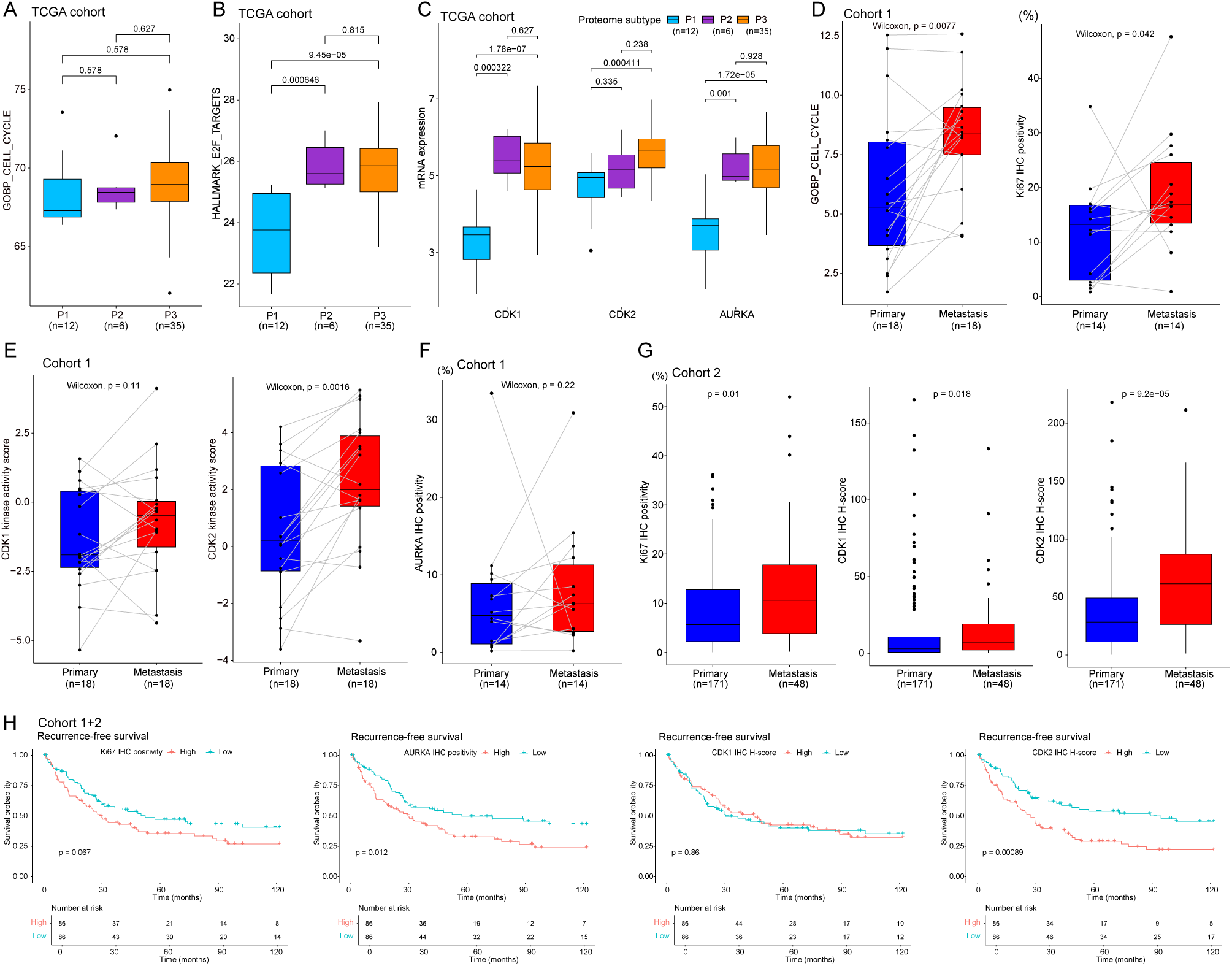
Cell cycle feature differences in the TCGA STLMS transcriptome data and comparison of metastatic samples in cohort 1 against matched primary samples, related to Figure 3. (A) Boxplot showing cell cycle score differences by proteome subtypes in the TCGA STLMS cohort. (B) Boxplot of “E2F targets” scores of ssGSEA by subtypes in the TCGA STLMS cohort. P2/P3 show significantly higher scores than P1. (C) Boxplot of mRNA expression of CDK1, CDK2, and AURKA in the TCGA cohort. P2/P3 show significantly higher expressions of CDK1 and AURKA than P1. CDK2 mRNA expression in P3 is higher than in P1. (D) Boxplot on the left side showing cell cycle score differences between 18 paired primary and metastatic samples in cohort 1. Boxplot on the right side showing Ki67-positive tumor cell frequency based on IHC by subtypes (14 paired samples) in cohort 1. (E) Boxplot showing kinase activity scores of CDK1/2 between primary and metastatic samples. 18 paired samples were analyzed from cohort 1. (F) Boxplot of AURKA IHC scores. 14 paired samples were available for assessment. (G) Boxplot of Ki67, CDK1, and CDK2 IHC scores between primary and metastasis samples in cohort 2, confirming metastatic samples show significant increases of these markers. (H) Kaplan-Meier analysis of recurrence-free survival in combined cohorts 1 and 2, stratified by Ki67, AURKA median positivities, CDK1/2 IHC H-score median values. Patients with high AURKA positivity or CDK2-high H-scores show poorer outcomes, concordant with poor survival outcome of the P2/P3 subtypes (cell cycle-enriched subtypes).

**Figure S4.**
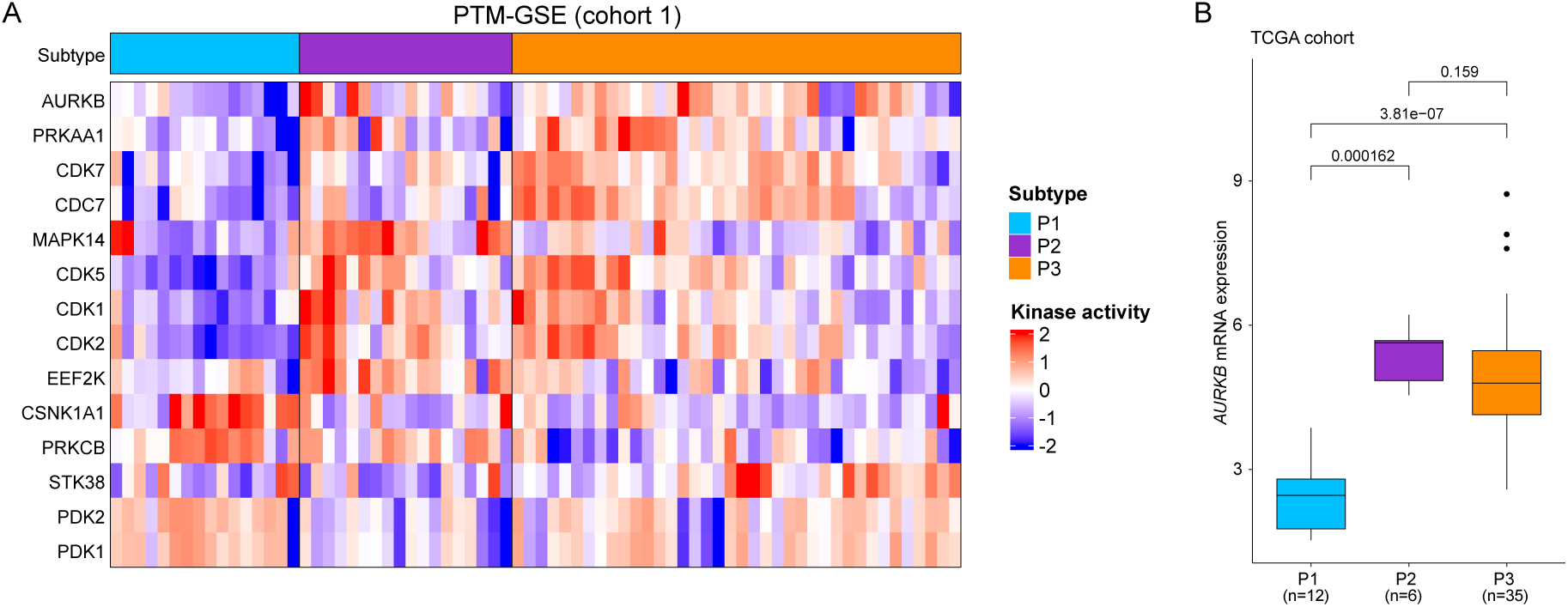
Phosphorylation landscape of cohort 1 and validation of findings in the independent cohort 2 and the external TCGA STLMS cohort, related to Figure 4. (A) A heatmap of PTM-GSE scores. Terms with significant differences between subtypes (ANOVA q<0.01) are shown. AURKB activity is high in P2/P3, consistent with the RoKAI results. (B) Boxplot of AURKB mRNA expression by subtypes in the TCGA cohort. AURKB mRNA expression is significantly higher in P2/P3 compared to P1.

**Figure S5.**
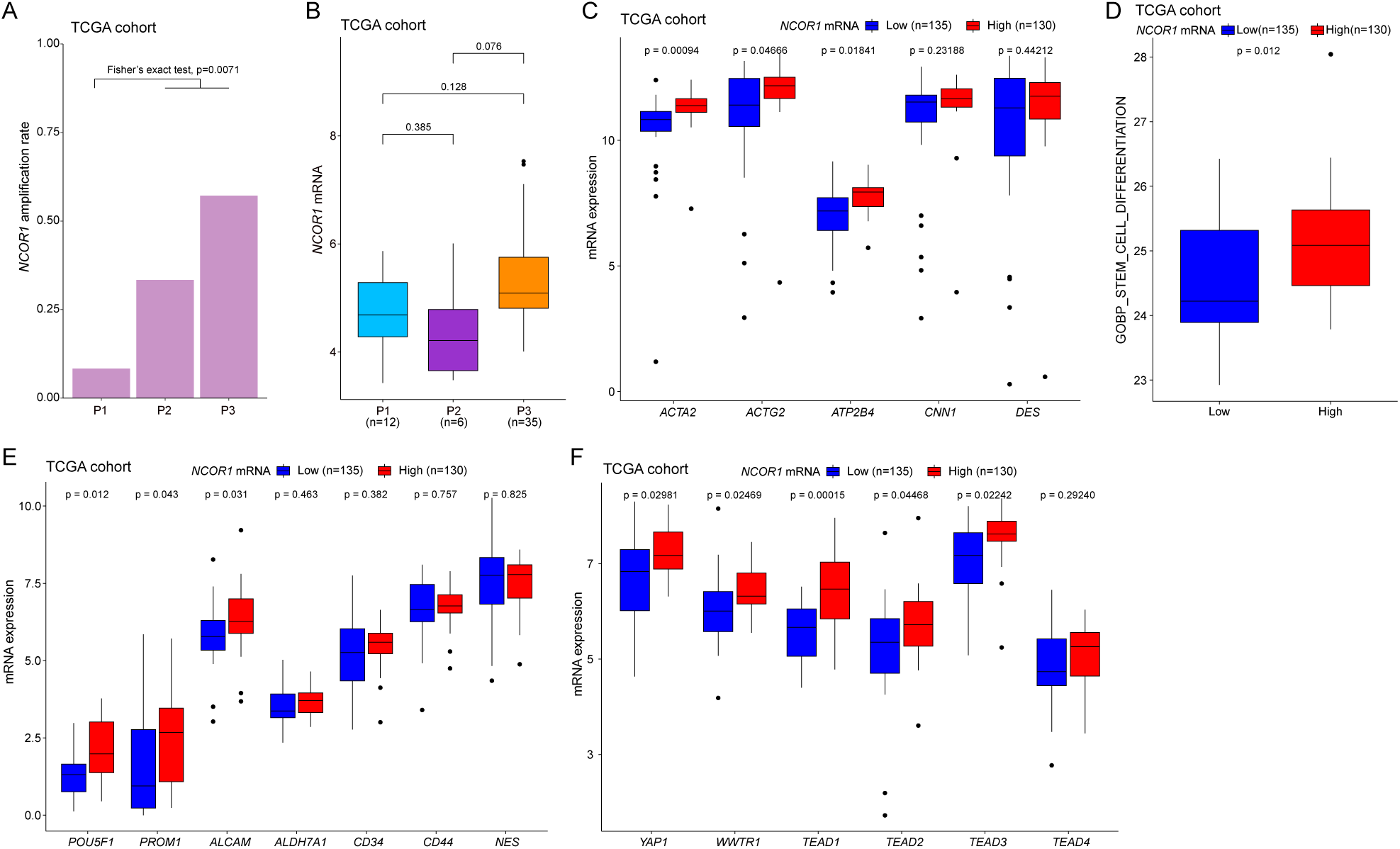
NCOR1 characteristics in the external TCGA STLMS dataset, related to Figure 5. (A) NCOR1 amplification shows higher frequency in P2/P3 compared to P1. (B) NCOR1 mRNA expression in P3 trends higher than in other subtypes, but not statistically significant. (C) mRNA expression of smooth muscle markers shows higher levels in the NCOR1-high group. (D) Stem cell differentiation scores (GOBP ssGSEA scores) are significantly higher in the NCOR1-high group. (E) mRNA expression of POU5F1, PROM1 (CD133), and ALCAM are significantly higher in the NCOR1-high group. (F) Boxplots of YAP1/TAZ (encoded by WWTR1) related molecules. mRNA expression of YAP1, WWTR1, and TEAD1/2/3 is significantly higher in the NCOR1-high group.

**Figure S6.**
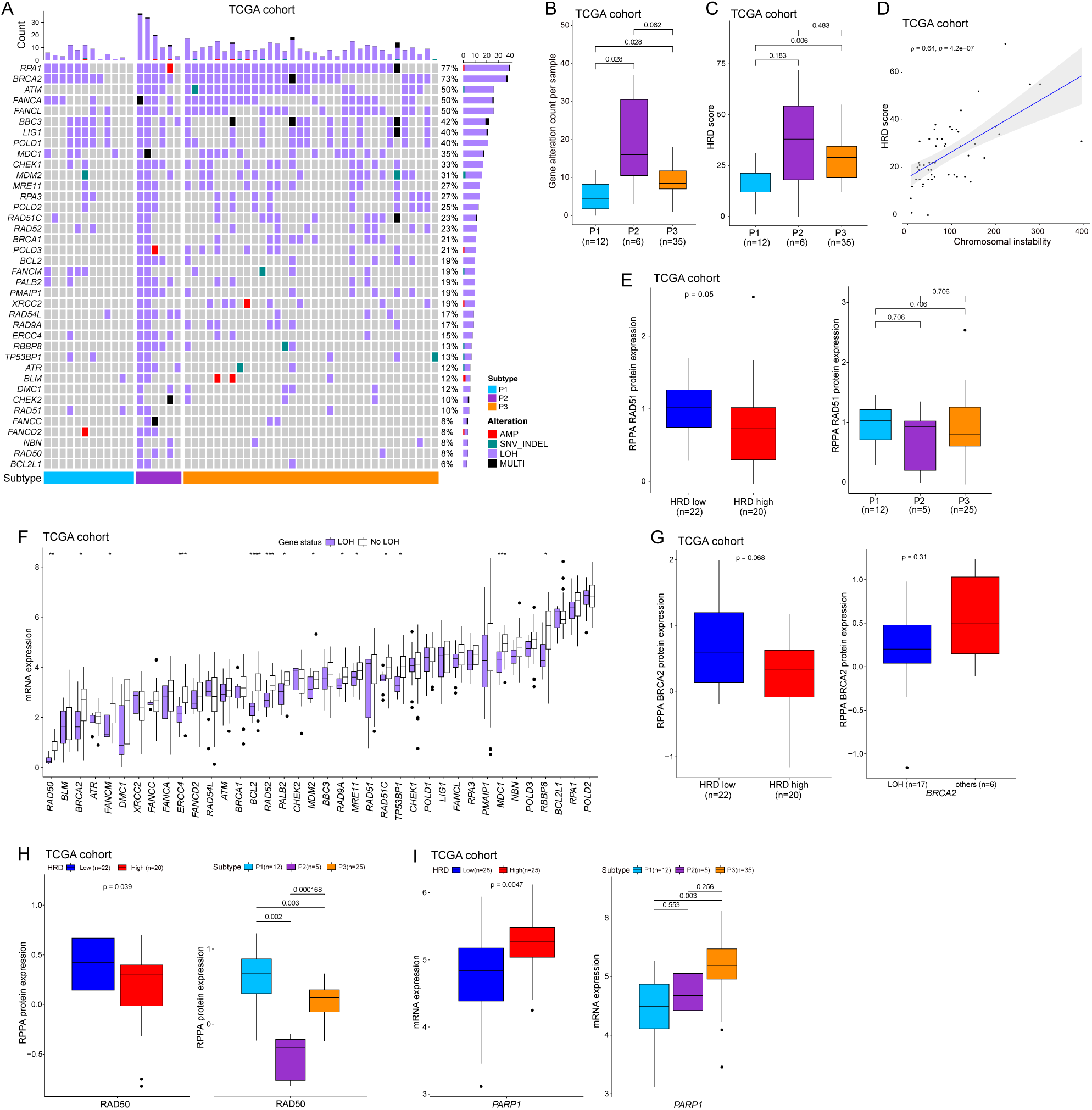
HRR pathway in validation cohort 2 and the external TCGA cohort, related to Figure 6. (A) Oncoprint of HRR pathway coponents showing frequent LOH events in the TCGA STLMS cohort. (B) Boxplot of genomic alteration counts in the HRR pathway per sample showing significantly higher occurrence in P2/P3 in the TCGA cohort. (C) The HRD score is higher in P2/P3 compared to P1 in the TCGA cohort. (D) The HRD score positively correlates with CIN in the TCGA cohort. (E) The RAD51 protein expression is lower in the HRD-low group and P2/P3 vs. P1, but not significant in the TCGA cohort. (F) Boxplots of mRNA expression between LOH and non-LOH groups for each HRR gene. All 38 HRR genes are shown. Most genes, including BRCA2, show lower expression in the LOH group compared to the non-LOH group. Note: all genes with statistical significance show expressional decrease in the LOH group in the TCGA cohort. (G) Boxplots of BRCA2 protein expression between HRD-low and HRD-high groups and BRCA2 LOH and non-LOH groups. BRCA2 protein expression shows the same trend as in cohort 1. (H) Boxplots of RAD50 protein expression between HRD-low and HRD-high groups and by proteome subtypes. RAD50 protein expression is significantly downregulated in the HRD-low group and in P2/P3 subtypes. (I) Boxplot of PARP1 mRNA expression between HRD-low and HRD-high groups and by proteome subtypes. PARP1 mRNA expression is significantly higher in the HRD-high group and the P3 subtype.

**Figure S7.**
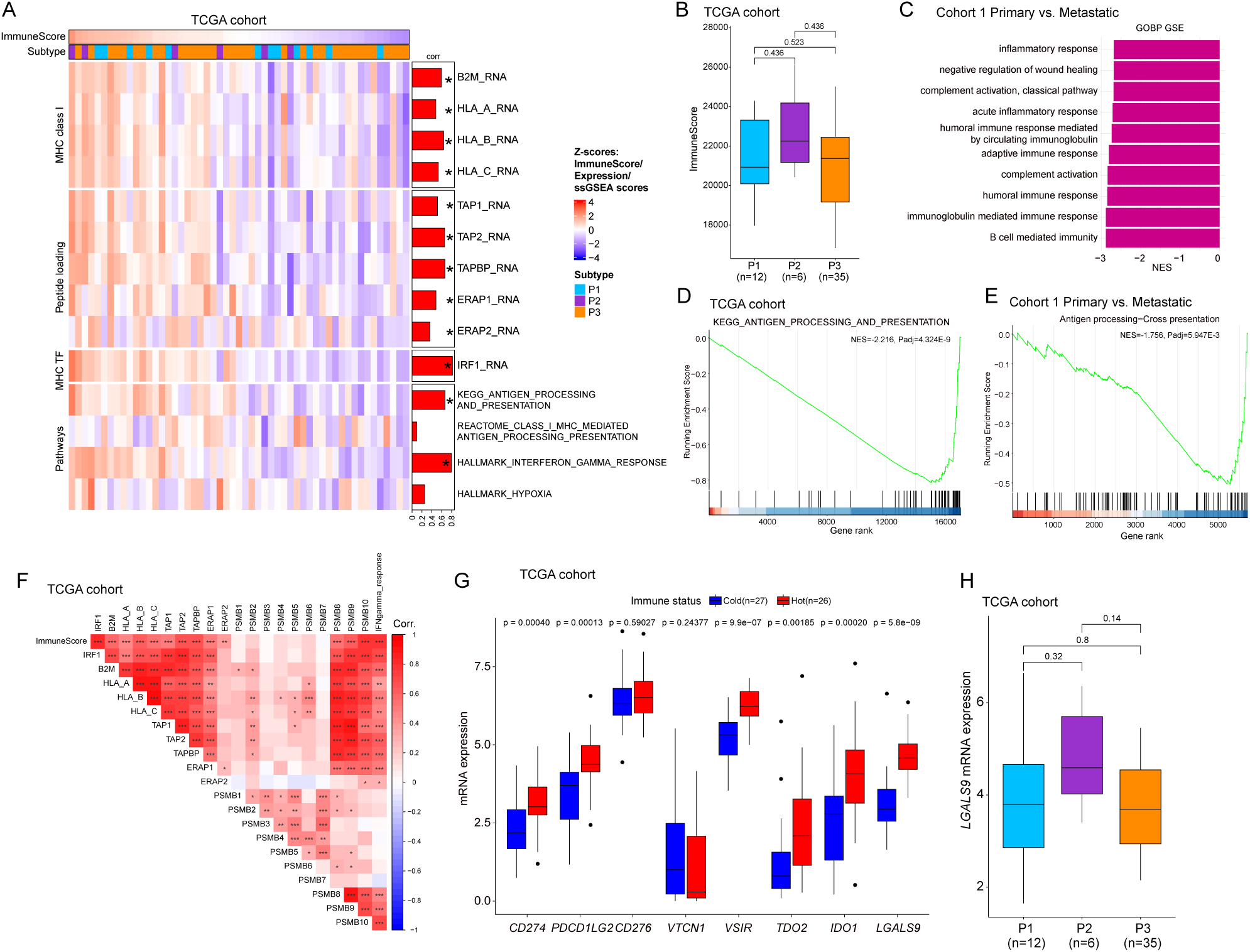
Tumor microenvironment analysis of the external TCGA STLMS dataset and comparison of primary vs. metastatic lesions, related to Figure 7. (A) A heatmap with immune signatures of the TCGA dataset. Immune-cold tumors show low expression of MHC class I molecules and peptide loading molecules, confirming findings in cohort 1. Bars on the right side of the heatmap denote Spearman’s correlation coefficients between ImmuneScores and each gene/signature. (B) Boxplot showing ImmuneScores by inferred proteome subtypes. Although not statistically significant, P3 shows lower scores compared to other subtypes. (C) Top 10 significantly downregulated GOBP terms. GSEA of differential expression between primary and metastatic tumors in cohort 1 shows significant downregulation of immune-related pathways in metastatic tumors compared to primary tumors. (D) GSE plot of KEGG pathway analyses showing significant downregulation of antigen processing pathway coponents in immune-cold tumors compared to immune-hot tumors in the TCGA dataset. (E) GSE plot of Reactome pathway analyses showing significant downregulation of antigen processing pathway components in metastatic tumors compared to primary tumors in cohort 1. (F) A correlation plot (Spearman’s correlation coefficients) including the ImmuneScore, IRF1, and antigen processing machinery protein expression in the TCGA cohort. *, p<0.05; **, p<0.01; ***, p<0.001. Most genes and ImmuneScore show positive correlation with each other. IFNgamma_response scores are ssGSEA scores of the “Hallmark_Interferon_Gamma_Reponse” term. (G) mRNA expressions of immunosuppressive genes in the TCGA cohort. All genes except CD276 and VTCN1 show significantly higher expression in immune-hot tumors compared to immune-cold tumors. (H) Boxplot of LGALS9 mRNA expression by subtypes in the TCGA cohort. P2 shows higher expression of LGALS9 compared to P1/P3, which is consistent with the findings in cohort 1.

## References

1. Soft Tissue and Bone Tumours. (2020). WHO Classification of Tumours, 5th edition Edition (IARC Publications).

2. Devaud, N., Vornicova, O., Abdul Razak, A.R., Khalili, K., Demicco, E.G., Mitric, C., Bernardini, M.Q., and Gladdy, R.A. (2022). Leiomyosarcoma: Current Clinical Management and Future Horizons. Surg Oncol Clin N Am 31, 527–546. 10.1016/j.soc.2022.03.011.

3. Serrano, C., and George, S. (2013). Leiomyosarcoma. Hematol Oncol Clin North Am 27, 957–974. 10.1016/j.hoc.2013.07.002.

4. Acem, I., van Houdt, W.J., Grünhagen, D.J., van der Graaf, W.T.A., Rueten-Budde, A.J., Gelderblom, H., Verhoef, C., and van de Sande, M.A.J. (2022). The role of perioperative chemotherapy in primary high-grade extremity soft tissue sarcoma: a risk-stratified analysis using PERSARC. European journal of cancer (Oxford, England : 1990) 165, 71-80. 10.1016/j.ejca.2022.01.013.

5. Gronchi, A., Miah, A.B., Dei Tos, A.P., Abecassis, N., Bajpai, J., Bauer, S., Biagini, R., Bielack, S., Blay, J.Y., Bolle, S., et al. (2021). Soft tissue and visceral sarcomas: ESMO-EURACAN-GENTURIS Clinical Practice Guidelines for diagnosis, treatment and follow-up(⋆). Ann Oncol 32, 1348–1365. 10.1016/j.annonc.2021.07.006.

6. Pisters, P.W., Leung, D.H., Woodruff, J., Shi, W., and Brennan, M.F. (1996). Analysis of prognostic factors in 1,041 patients with localized soft tissue sarcomas of the extremities. Journal of clinical oncology : official journal of the American Society of Clinical Oncology 14, 1679–1689. 10.1200/jco.1996.14.5.1679.

7. Lewis, J.J., Leung, D., Heslin, M., Woodruff, J.M., and Brennan, M.F. (1997). Association of local recurrence with subsequent survival in extremity soft tissue sarcoma. Journal of clinical oncology : official journal of the American Society of Clinical Oncology 15, 646–652. 10.1200/jco.1997.15.2.646.

8. Trovik, C.S., Bauer, H.C., Alvegård, T.A., Anderson, H., Blomqvist, C., Berlin, O., Gustafson, P., Saeter, G., and Wallöe, A. (2000). Surgical margins, local recurrence and metastasis in soft tissue sarcomas: 559 surgically-treated patients from the Scandinavian Sarcoma Group Register. European journal of cancer (Oxford, England : 1990) 36, 710–716. 10.1016/s0959-8049(99)00287-7.

9. Linch, M., Miah, A.B., Thway, K., Judson, I.R., and Benson, C. (2014). Systemic treatment of soft-tissue sarcoma-gold standard and novel therapies. Nat Rev Clin Oncol 11, 187–202. 10.1038/nrclinonc.2014.26.

10. Savina, M., Le Cesne, A., Blay, J.Y., Ray-Coquard, I., Mir, O., Toulmonde, M., Cousin, S., Terrier, P., Ranchere-Vince, D., Meeus, P., et al. (2017). Patterns of care and outcomes of patients with METAstatic soft tissue SARComa in a real-life setting: the METASARC observational study. BMC Med 15, 78. 10.1186/s12916-017-0831-7.

11. Chibon, F., Lagarde, P., Salas, S., Pérot, G., Brouste, V., Tirode, F., Lucchesi, C., de Reynies, A., Kauffmann, A., Bui, B., et al. (2010). Validated prediction of clinical outcome in sarcomas and multiple types of cancer on the basis of a gene expression signature related to genome complexity. Nature medicine 16, 781–787. 10.1038/nm.2174.

12. TCGA (2017). Comprehensive and Integrated Genomic Characterization of Adult Soft Tissue Sarcomas. Cell 171, 950–965.e928. 10.1016/j.cell.2017.10.014.

13. Nacev, B.A., Sanchez-Vega, F., Smith, S.A., Antonescu, C.R., Rosenbaum, E., Shi, H., Tang, C., Socci, N.D., Rana, S., Gularte-Merida, R., et al. (2022). Clinical sequencing of soft tissue and bone sarcomas delineates diverse genomic landscapes and potential therapeutic targets. Nat Commun 13, 3405. 10.1038/s41467-022-30453-x.

14. Italiano, A., Kind, M., Stoeckle, E., Jones, N., Coindre, J.M., and Bui, B. (2011). Temsirolimus in advanced leiomyosarcomas: patterns of response and correlation with the activation of the mammalian target of rapamycin pathway. Anticancer Drugs 22, 463–467. 10.1097/CAD.0b013e3283442074.

15. Schwartz, G.K., Tap, W.D., Qin, L.X., Livingston, M.B., Undevia, S.D., Chmielowski, B., Agulnik, M., Schuetze, S.M., Reed, D.R., Okuno, S.H., et al. (2013). Cixutumumab and temsirolimus for patients with bone and soft-tissue sarcoma: a multicentre, open-label, phase 2 trial. The Lancet. Oncology 14, 371–382. 10.1016/s1470-2045(13)70049-4.

16. Burns, J., Wilding, C.P., R, L.J., and P, H.H. (2020). Proteomic research in sarcomas - current status and future opportunities. Semin Cancer Biol 61, 56–70. 10.1016/j.semcancer.2019.11.003.

17. Gillette, M.A., Satpathy, S., Cao, S., Dhanasekaran, S.M., Vasaikar, S.V., Krug, K., Petralia, F., Li, Y., Liang, W.W., Reva, B., et al. (2020). Proteogenomic Characterization Reveals Therapeutic Vulnerabilities in Lung Adenocarcinoma. Cell 182, 200–225.e235. 10.1016/j.cell.2020.06.013.

18. Zhang, B., Wang, J., Wang, X., Zhu, J., Liu, Q., Shi, Z., Chambers, M.C., Zimmerman, L.J., Shaddox, K.F., Kim, S., et al. (2014). Proteogenomic characterization of human colon and rectal cancer. Nature 513, 382–387. 10.1038/nature13438.

19. Burns, J., Wilding, C.P., Krasny, L., Zhu, X., Chadha, M., Tam, Y.B., Ps, H., Mahalingam, A.H., Lee, A.T.J., Arthur, A., et al. (2023). The proteomic landscape of soft tissue sarcomas. Nat Commun 14, 3834. 10.1038/s41467-023-39486-2.

20. Tang, S., Wang, Y., Luo, R., Fang, R., Liu, Y., Xiang, H., Ran, P., Tong, Y., Sun, M., Tan, S., et al. (2024). Proteomic characterization identifies clinically relevant subgroups of soft tissue sarcoma. Nat Commun 15, 1381. 10.1038/s41467-024-45306-y.

21. AACR Project GENIE: Powering Precision Medicine through an International Consortium. (2017). Cancer Discov 7, 818–831. 10.1158/2159-8290.Cd-17-0151.

22. Ucar, D., and Lin, D.I. (2015). Amplification of the bromodomain-containing protein 4 gene in ovarian high-grade serous carcinoma is associated with worse prognosis and survival. Mol Clin Oncol 3, 1291–1294. 10.3892/mco.2015.622.

23. Brahmi, M., Lesluyes, T., Dufresne, A., Toulmonde, M., Italiano, A., Mir, O., Le Cesne, A., Valentin, T., Chevreau, C., Bonvalot, S., et al. (2021). Expression and prognostic significance of PDGF ligands and receptors across soft tissue sarcomas. ESMO Open 6, 100037. 10.1016/j.esmoop.2020.100037.

24. Peng, L., Li, J., Wu, J., Xu, B., Wang, Z., Giamas, G., Stebbing, J., and Yu, Z. (2021). A Pan-Cancer Analysis of SMARCA4 Alterations in Human Cancers. Front Immunol 12, 762598. 10.3389/fimmu.2021.762598.

25. Liang, J., Li, B., Yuan, L., and Ye, Z. (2015). Prognostic value of IGF-1R expression in bone and soft tissue sarcomas: a meta-analysis. Onco Targets Ther 8, 1949–1955. 10.2147/ott.S88293.

26. Li, Y., Li, Y., Liu, Y., Xie, P., Li, F., and Li, G. (2014). PAX6, a novel target of microRNA-7, promotes cellular proliferation and invasion in human colorectal cancer cells. Dig Dis Sci 59, 598–606. 10.1007/s10620-013-2929-x.

27. Ooki, A., Dinalankara, W., Marchionni, L., Tsay, J.J., Goparaju, C., Maleki, Z., Rom, W.N., Pass, H.I., and Hoque, M.O. (2018). Epigenetically regulated PAX6 drives cancer cells toward a stem-like state via GLI-SOX2 signaling axis in lung adenocarcinoma. Oncogene 37, 5967–5981. 10.1038/s41388-018-0373-2.

28. Jing, N., Du, X., Liang, Y., Tao, Z., Bao, S., Xiao, H., Dong, B., Gao, W.Q., and Fang, Y.X. (2024). PAX6 promotes neuroendocrine phenotypes of prostate cancer via enhancing MET/STAT5A-mediated chromatin accessibility. J Exp Clin Cancer Res 43, 144. 10.1186/s13046-024-03064-1.

29. Ballotti, R., Cheli, Y., and Bertolotto, C. (2020). The complex relationship between MITF and the immune system: a Melanoma ImmunoTherapy (response) Factor? Molecular cancer 19, 170. 10.1186/s12943-020-01290-7.

30. Du, R., Huang, C., Liu, K., Li, X., and Dong, Z. (2021). Targeting AURKA in Cancer: molecular mechanisms and opportunities for Cancer therapy. Molecular cancer 20, 15. 10.1186/s12943-020-01305-3.

31. Yilmaz, S., Ayati, M., Schlatzer, D., Cicek, A.E., Chance, M.R., and Koyuturk, M. (2021). Robust inference of kinase activity using functional networks. Nat Commun 12, 1177. 10.1038/s41467-021-21211-6.

32. Jaenisch, R., and Bird, A. (2003). Epigenetic regulation of gene expression: how the genome integrates intrinsic and environmental signals. Nature genetics 33 *Suppl*, 245-254. 10.1038/ng1089.

33. Li, X., and Heyer, W.D. (2008). Homologous recombination in DNA repair and DNA damage tolerance. Cell Res 18, 99–113. 10.1038/cr.2008.1.

34. Lord, C.J., and Ashworth, A. (2016). BRCAness revisited. Nature reviews. Cancer 16, 110–120. 10.1038/nrc.2015.21.

35. Turajlic, S., and Swanton, C. (2016). Metastasis as an evolutionary process. Science (New York, N.Y.) 352, 169–175. 10.1126/science.aaf2784.

36. Bakhoum, S.F., Ngo, B., Laughney, A.M., Cavallo, J.A., Murphy, C.J., Ly, P., Shah, P., Sriram, R.K., Watkins, T.B.K., Taunk, N.K., et al. (2018). Chromosomal instability drives metastasis through a cytosolic DNA response. Nature 553, 467–472. 10.1038/nature25432.

37. Planas-Paz, L., Pliego-Mendieta, A., Hagedorn, C., Aguilera-Garcia, D., Haberecker, M., Arnold, F., Herzog, M., Bankel, L., Guggenberger, R., Steiner, S., et al. (2023). Unravelling homologous recombination repair deficiency and therapeutic opportunities in soft tissue and bone sarcoma. EMBO Mol Med 15, e16863. 10.15252/emmm.202216863.

38. Baumann, P., and West, S.C. (1998). Role of the human RAD51 protein in homologous recombination and double-stranded-break repair. Trends Biochem Sci 23, 247–251. 10.1016/s0968-0004(98)01232-8.

39. Mansour, W.Y., Rhein, T., and Dahm-Daphi, J. (2010). The alternative end-joining pathway for repair of DNA double-strand breaks requires PARP1 but is not dependent upon microhomologies. Nucleic Acids Res 38, 6065–6077. 10.1093/nar/gkq387.

40. Yoshihara, K., Shahmoradgoli, M., Martínez, E., Vegesna, R., Kim, H., Torres-Garcia, W., Treviño, V., Shen, H., Laird, P.W., Levine, D.A., et al. (2013). Inferring tumour purity and stromal and immune cell admixture from expression data. Nat Commun 4, 2612. 10.1038/ncomms3612.

41. Zhou, F. (2009). Molecular mechanisms of IFN-gamma to up-regulate MHC class I antigen processing and presentation. Int Rev Immunol 28, 239–260. 10.1080/08830180902978120.

42. Der, S.D., Zhou, A., Williams, B.R., and Silverman, R.H. (1998). Identification of genes differentially regulated by interferon alpha, beta, or gamma using oligonucleotide arrays. Proceedings of the National Academy of Sciences of the United States of America 95, 15623–15628. 10.1073/pnas.95.26.15623.

43. Kikutake, C., and Suyama, M. (2023). Pan-cancer analysis of whole-genome doubling and its association with patient prognosis. BMC cancer 23, 619. 10.1186/s12885-023-11132-6.

44. Baker, T.M., Lai, S., Lynch, A.R., Lesluyes, T., Yan, H., Ogilvie, H.A., Verfaillie, A., Dentro, S., Bowes, A.L., Pillay, N., et al. (2024). The History of Chromosomal Instability in Genome-Doubled Tumors. Cancer Discov 14, 1810–1822. 10.1158/2159-8290.Cd-23-1249.

45. Nikhil, K., and Shah, K. (2024). The significant others of aurora kinase a in cancer: combination is the key. Biomark Res 12, 109. 10.1186/s40364-024-00651-4.

46. Jamalzadeh, S., Dai, J., Lavikka, K., Li, Y., Jiang, J., Huhtinen, K., Virtanen, A., Oikkonen, J., Hietanen, S., Hynninen, J., et al. (2024). Genome-wide quantification of copy-number aberration impact on gene expression in ovarian high-grade serous carcinoma. BMC cancer 24, 173. 10.1186/s12885-024-11895-6.

47. Yamamoto, H., Williams, E.G., Mouchiroud, L., Cantó, C., Fan, W., Downes, M., Héligon, C., Barish, G.D., Desvergne, B., Evans, R.M., et al. (2011). NCoR1 is a conserved physiological modulator of muscle mass and oxidative function. Cell 147, 827–839. 10.1016/j.cell.2011.10.017.

48. Kohn, E.C., Lee, J.M., and Ivy, S.P. (2017). The HRD Decision-Which PARP Inhibitor to Use for Whom and When. Clin Cancer Res 23, 7155–7157. 10.1158/1078-0432.Ccr-17-2186.

49. Dall, G., Vandenberg, C.J., Nesic, K., Ratnayake, G., Zhu, W., Vissers, J.H.A., Bedő, J., Penington, J., Wakefield, M.J., Kee, D., et al. (2023). Targeting homologous recombination deficiency in uterine leiomyosarcoma. J Exp Clin Cancer Res 42, 112. 10.1186/s13046-023-02687-0.

50. Slade, D., and Loizou, J.I. (2023). Leveraging homologous recombination repair deficiency in sarcoma. EMBO Mol Med 15, e17453. 10.15252/emmm.202317453.

51. Cheng, D.T., Mitchell, T.N., Zehir, A., Shah, R.H., Benayed, R., Syed, A., Chandramohan, R., Liu, Z.Y., Won, H.H., Scott, S.N., et al. (2015). Memorial Sloan Kettering-Integrated Mutation Profiling of Actionable Cancer Targets (MSK-IMPACT): A Hybridization Capture-Based Next-Generation Sequencing Clinical Assay for Solid Tumor Molecular Oncology. The Journal of molecular diagnostics : JMD 17, 251–264. 10.1016/j.jmoldx.2014.12.006.

52. Li, H., and Durbin, R. (2009). Fast and accurate short read alignment with Burrows-Wheeler transform. Bioinformatics (Oxford, England) 25, 1754–1760. 10.1093/bioinformatics/btp324.

53. DePristo, M.A., Banks, E., Poplin, R., Garimella, K.V., Maguire, J.R., Hartl, C., Philippakis, A.A., del Angel, G., Rivas, M.A., Hanna, M., et al. (2011). A framework for variation discovery and genotyping using next-generation DNA sequencing data. Nature genetics 43, 491–498. 10.1038/ng.806.

54. Mose, L.E., Wilkerson, M.D., Hayes, D.N., Perou, C.M., and Parker, J.S. (2014). ABRA: improved coding indel detection via assembly-based realignment. Bioinformatics (Oxford, England) 30, 2813–2815. 10.1093/bioinformatics/btu376.

55. Cibulskis, K., Lawrence, M.S., Carter, S.L., Sivachenko, A., Jaffe, D., Sougnez, C., Gabriel, S., Meyerson, M., Lander, E.S., and Getz, G. (2013). Sensitive detection of somatic point mutations in impure and heterogeneous cancer samples. Nature biotechnology 31, 213–219. 10.1038/nbt.2514.

56. Ye, K., Schulz, M.H., Long, Q., Apweiler, R., and Ning, Z. (2009). Pindel: a pattern growth approach to detect break points of large deletions and medium sized insertions from paired-end short reads. Bioinformatics (Oxford, England) 25, 2865–2871. 10.1093/bioinformatics/btp394.

57. Lai, Z., Markovets, A., Ahdesmaki, M., Chapman, B., Hofmann, O., McEwen, R., Johnson, J., Dougherty, B., Barrett, J.C., and Dry, J.R. (2016). VarDict: a novel and versatile variant caller for next-generation sequencing in cancer research. Nucleic Acids Res 44, e108. 10.1093/nar/gkw227.

58. Shen, R., and Seshan, V.E. (2016). FACETS: allele-specific copy number and clonal heterogeneity analysis tool for high-throughput DNA sequencing. Nucleic Acids Res 44, e131. 10.1093/nar/gkw520.

59. Bielski, C.M., Zehir, A., Penson, A.V., Donoghue, M.T.A., Chatila, W., Armenia, J., Chang, M.T., Schram, A.M., Jonsson, P., Bandlamudi, C., et al. (2018). Genome doubling shapes the evolution and prognosis of advanced cancers. Nature genetics 50, 1189–1195. 10.1038/s41588-018-0165-1.

60. Colaprico, A., Silva, T.C., Olsen, C., Garofano, L., Cava, C., Garolini, D., Sabedot, T.S., Malta, T.M., Pagnotta, S.M., Castiglioni, I., et al. (2016). TCGAbiolinks: an R/Bioconductor package for integrative analysis of TCGA data. Nucleic Acids Res 44, e71. 10.1093/nar/gkv1507.

61. Sztupinszki, Z., Diossy, M., Krzystanek, M., Reiniger, L., Csabai, I., Favero, F., Birkbak, N.J., Eklund, A.C., Syed, A., and Szallasi, Z. (2018). Migrating the SNP array-based homologous recombination deficiency measures to next generation sequencing data of breast cancer. NPJ Breast Cancer 4, 16. 10.1038/s41523-018-0066-6.

62. Shi, Z., Chen, B., Han, X., Gu, W., Liang, S., and Wu, L. (2023). Genomic and molecular landscape of homologous recombination deficiency across multiple cancer types. Scientific reports 13, 8899. 10.1038/s41598-023-35092-w.

63. Mermel, C.H., Schumacher, S.E., Hill, B., Meyerson, M.L., Beroukhim, R., and Getz, G. (2011). GISTIC2.0 facilitates sensitive and confident localization of the targets of focal somatic copy-number alteration in human cancers. Genome Biol 12, R41. 10.1186/gb-2011-12-4-r41.

64. Dobin, A., Davis, C.A., Schlesinger, F., Drenkow, J., Zaleski, C., Jha, S., Batut, P., Chaisson, M., and Gingeras, T.R. (2013). STAR: ultrafast universal RNA-seq aligner. Bioinformatics (Oxford, England) 29, 15–21. 10.1093/bioinformatics/bts635.

65. Li, B., and Dewey, C.N. (2011). RSEM: accurate transcript quantification from RNA-Seq data with or without a reference genome. BMC bioinformatics 12, 323. 10.1186/1471-2105-12-323.

66. McAlister, G.C., Nusinow, D.P., Jedrychowski, M.P., Wuhr, M., Huttlin, E.L., Erickson, B.K., Rad, R., Haas, W., and Gygi, S.P. (2014). MultiNotch MS3 Enables Accurate, Sensitive, and Multiplexed Detection of Differential Expression across Cancer Cell Line Proteomes. Analytical Chemistry 86, 7150–7158. 10.1021/ac502040v.

67. Navarrete-Perea, J., Yu, Q., Gygi, S.P., and Paulo, J.A. (2018). Streamlined Tandem Mass Tag (SL-TMT) Protocol: An Efficient Strategy for Quantitative (Phospho)proteome Profiling Using Tandem Mass Tag-Synchronous Precursor Selection-MS3. J Proteome Res 17, 2226–2236. 10.1021/acs.jproteome.8b00217.

68. Liu, H., Sadygov, R.G., and Yates, J.R., 3rd (2004). A model for random sampling and estimation of relative protein abundance in shotgun proteomics. Analytical chemistry 76, 4193–4201. 10.1021/ac0498563.

69. Plubell, D.L., Wilmarth, P.A., Zhao, Y., Fenton, A.M., Minnier, J., Reddy, A.P., Klimek, J., Yang, X., David, L.L., and Pamir, N. (2017). Extended Multiplexing of Tandem Mass Tags (TMT) Labeling Reveals Age and High Fat Diet Specific Proteome Changes in Mouse Epididymal Adipose Tissue. Mol Cell Proteomics 16, 873–890. 10.1074/mcp.M116.065524.

70. Robinson, M.D., McCarthy, D.J., and Smyth, G.K. (2010). edgeR: a Bioconductor package for differential expression analysis of digital gene expression data. Bioinformatics (Oxford, England) 26, 139–140. 10.1093/bioinformatics/btp616.

71. Gaujoux, R., and Seoighe, C. (2010). A flexible R package for nonnegative matrix factorization. BMC bioinformatics 11, 367. 10.1186/1471-2105-11-367.

72. Kim, H., and Park, H. (2007). Sparse non-negative matrix factorizations via alternating non-negativity-constrained least squares for microarray data analysis. Bioinformatics (Oxford, England) 23, 1495–1502. 10.1093/bioinformatics/btm134.

73. Yu, G., Wang, L.G., Han, Y., and He, Q.Y. (2012). clusterProfiler: an R package for comparing biological themes among gene clusters. Omics 16, 284–287. 10.1089/omi.2011.0118.

74. Barbie, D.A., Tamayo, P., Boehm, J.S., Kim, S.Y., Moody, S.E., Dunn, I.F., Schinzel, A.C., Sandy, P., Meylan, E., Scholl, C., et al. (2009). Systematic RNA interference reveals that oncogenic KRAS-driven cancers require TBK1. Nature 462, 108–112. 10.1038/nature08460.

75. Hornbeck, P.V., Zhang, B., Murray, B., Kornhauser, J.M., Latham, V., and Skrzypek, E. (2015). PhosphoSitePlus, 2014: mutations, PTMs and recalibrations. Nucleic Acids Res 43, D512-520. 10.1093/nar/gku1267.

76. Castro, M.A., de Santiago, I., Campbell, T.M., Vaughn, C., Hickey, T.E., Ross, E., Tilley, W.D., Markowetz, F., Ponder, B.A., and Meyer, K.B. (2016). Regulators of genetic risk of breast cancer identified by integrative network analysis. Nature genetics 48, 12–21. 10.1038/ng.3458.

77. Fletcher, M.N., Castro, M.A., Wang, X., de Santiago, I., O’Reilly, M., Chin, S.F., Rueda, O.M., Caldas, C., Ponder, B.A., Markowetz, F., and Meyer, K.B. (2013). Master regulators of FGFR2 signalling and breast cancer risk. Nat Commun 4, 2464. 10.1038/ncomms3464.

78. Robertson, A.G., Kim, J., Al-Ahmadie, H., Bellmunt, J., Guo, G., Cherniack, A.D., Hinoue, T., Laird, P.W., Hoadley, K.A., Akbani, R., et al. (2017). Comprehensive Molecular Characterization of Muscle-Invasive Bladder Cancer. Cell 171, 540–556.e525. 10.1016/j.cell.2017.09.007.

79. Lambert, S.A., Jolma, A., Campitelli, L.F., Das, P.K., Yin, Y., Albu, M., Chen, X., Taipale, J., Hughes, T.R., and Weirauch, M.T. (2018). The Human Transcription Factors. Cell 172, 650–665. 10.1016/j.cell.2018.01.029.

80. Margolin, A.A., Nemenman, I., Basso, K., Wiggins, C., Stolovitzky, G., Dalla Favera, R., and Califano, A. (2006). ARACNE: an algorithm for the reconstruction of gene regulatory networks in a mammalian cellular context. BMC bioinformatics 7 *Suppl 1*, S7. 10.1186/1471-2105-7-s1-s7.

81. Carro, M.S., Lim, W.K., Alvarez, M.J., Bollo, R.J., Zhao, X., Snyder, E.Y., Sulman, E.P., Anne, S.L., Doetsch, F., Colman, H., et al. (2010). The transcriptional network for mesenchymal transformation of brain tumours. Nature 463, 318–325. 10.1038/nature08712.

82. Tsherniak, A., Vazquez, F., Montgomery, P.G., Weir, B.A., Kryukov, G., Cowley, G.S., Gill, S., Harrington, W.F., Pantel, S., Krill-Burger, J.M., et al. (2017). Defining a Cancer Dependency Map. Cell 170, 564–576.e516. 10.1016/j.cell.2017.06.010.

83. Kuhn M, W.H. (2020). Tidymodels: a collection of packages for modeling and machine learning using tidyverse principles.

